# Exploiting NDH-2 Vulnerability: Quinolines as Antitubercular Agents

**DOI:** 10.64898/2026.06.08.730781

**Authors:** Shashikanta Sau, Rohit Kumar, Arnab Roy, Puja Kumari Agnivesh, Pallavi Saha, Harshada Anil Bhalerao, Rajesh Sonti, Deepak Kumar Sharma, Nitin Pal Kalia

**Affiliations:** Department of Biological Sciences (Pharmacology and Toxicology), National Institute of Pharmaceutical Education and Research Hyderabad (NIPER-H), Telangana 500037, India; Department of Pharmaceutical Engg. and Tech., IIT-Banaras Hindu University, Varanasi, Uttar Pradesh 221005, India; Department of Pharmaceutical Analysis, National Institute of Pharmaceutical Education and Research Hyderabad (NIPER-H), Telangana 500037, India

**Keywords:** Electron transport chain (ETC), NADH dehydrogenase-II (NDH-2), Multi drug resistant (MDR), Oxygen consumption rate (OCR), Quinoline

## Abstract

*Mycobacterium tuberculosis* possesses a flexible metabolic system helping it to survive inside the host. The type II NADH dehydrogenase, composed of Ndh and NdhA, essential for bacilli, is a promising drug target. Based on ATP depletion values, two quinoline scaffolds were shortlisted after screening of a library of drug like molecules. Structurally, both **64-9C** and **64-9D** carry ester moieties at the 5– and 8-positions of the quinoline core, respectively. Ease to re-synthesise 64-9D resulted in synthesis of a focused library of compounds, with MIC values of 0.25–4 μg/mL, consistent with ATP depletion. These compounds exhibited bactericidal activity against non-replicating mycobacteria, and showed potent efficacy against multidrug-resistant isolates. Altered, intracellular NADH/NAD^+^ ratio and reduced respiration was indicative of oxidative phosphorylation inhibition. Inhibition of the purified recombinant NDH protein uncompetitively, SNPs in gene encoding NDH-2 for selected one step mutants and, molecular modelling of 4FQN and 2FQN validated NDH-2 as a target for these compounds. The derivative 2FQN exhibited dose-dependent bactericidal efficacy in mice, underscoring the potential of the series as a promising anti-tuberculosis candidates.

## Introduction

Tuberculosis (TB) caused by *Mtb* is only second to HIV in deaths due to infectious causes and the 13^th^ largest cause of death. In 2024, approximately 10.7 million cases of this illness were reported worldwide.(Global tuberculosis report 2025, 2025), (2024 Global tuberculosis report, 2024) The pulmonary tuberculosis is most common amongst different form of TB diseases. Active tuberculosis disease spreads through aerosols generated while breathing, singing, or coughing.(Field *et al*, 2018) The treatment for MDR-TB treatment is challenging due to use of multidrug regimen for a very long duration (18-24 months). Such a long treatment leads to poor compliance and has poor outcome with an estimated global success rate of 52%. Thus MDR-TB is a serious threat to the global human health.(Kalia *et al*, 2017a)

Conventional antibiotics that inhibit cell wall biosynthesis, such as β-lactams and isoniazid, are highly effective only against actively dividing bacilli but not effective against nonreplicating *Mycobacterium tuberculosis*, even when administered at concentrations up to 500-fold higher than their minimum inhibitory concentrations (MICs).(Koul *et al*, 2014) After a long gap of 40 years bedaquiline (BDQ) was FDA approved in 2012 for treatment of MDR-TB. But in 2014, soon after its usage in patients, the BDQ resistant TB strains emerged. Nonetheless, discovery of BDQ validated the Ox-Phos pathway as a valid target for anti-TB drug discovery and opened the avenues for exploiting components of electron transport chain (ETC) for identification of new chemical entities as anti-TB agents. The concept was additionally validated with the discovery of Telacebec (Q203), a potent investigational antibiotic developed by TB Alliance targeting the cytochrome bc1-*aa3* supercomplex of ETC.(Pethe *et al*, 2013) This discovery not only validated Cytochrome *bc1-aa3* complex as a vulnerable node for chemotherapeutic intervention but further emphasized the essentiality of Ox-Phos in *Mtb* survival. In continuation, the researchers from various scientific groups identified NADH dehydrogenases (NDH-2), succinate dehydrogenase (SDH), and cytochrome *bd* oxidase as viable drug targets.(Lu *et al*, 2018; Cook *et al*, 2017; Kalia *et al*, 2017a; Pecsi *et al*, 2014)

In *Mtb*, the electron transport chain (ETC) connects to the tricarboxylic acid (TCA) cycle through two major dehydrogenases: the proton-pumping type I NADH dehydrogenase and the non-proton-pumping type II NADHdehydrogenase.(Vilchèze *et al*, 2018) Electrons derived from these complexes are shuttled to the Cytochrome *bc1-aa_3_* complex via menaquinones and are subsequently transferred to molecular oxygen via the cytochrome bd oxidase, the terminal oxidase present in the ETC. Among these, NDH-2, a non-proton-pumping enzyme facilitating NADH oxidation and electron transfer to menaquinones, plays a central role in maintaining respiratory ATP synthesis. Given its vital role across drug-susceptible and drug-resistant strains, NDH-2 represents a promising target for the development of next-generation anti-mycobacterial agents.(Bhat *et al*, 2016) Notably, NDH-2 is absent in mitochondria, thereby offering a safer therapeutic window with potentially no host toxicity. Furthermore, its functional role under both replicating and non-replicating conditions underscores its essentiality during active and persistent phases of infection.(Saha *et al*, 2024b) Therefore, considering the importance of NDH-2 there is an urgent need to identify new chemical entities that inhibit the function of this enzyme.

This study was aimed on identification of new chemical class of *Mtb* NDH-2 inhibitors. Further, it is expected that the identified NDH-2 inhibitors may have synergy with BDQ and inhibitors of other components of ETC pathway with the hope of complete sterilization, shortening of treatment duration and minimum chances of emergence of resistant strains of *Mtb*.

## Results

### Whole cell primary screening of a library of drug like small molecules to identify new scaffolds targeting NDH-2

In order to identify whole cell active compounds, a commercially available library of 10,000 drug-like small molecules was screened at a single concentration of 10 µg/mL, keeping ATP level as a marker (Figure 2A).(Kalia *et al*, 2017a; Koul *et al*, 2014; Rao *et al*, 2008) Due to biosafety concerns, the initial screen was performed using *M. bovis* BCG. *M. bovis* share 99.9% similarity with *Mtb* at genomic level and is an excellent surrogate mycobacterium for drug screening. The screening was done in the absence and presence of Q203. This led to identification of two quinoline based small molecules i.e. quinolin-5-yl 2-fluorobenzoate (**64-9C**) & quinolin-8-yl 3-nitrobenzoate (64-9D), as they were able to deplete the intracellular ATP in both conditions. Compounds 64-9C and 64-9D were further subjected for growth inhibition assays for determination of MIC and ATP IC_50_ and *in-silico* binding approach evaluation (Figure 2B). Furthermore, we observed that the molecule 64-9D had a good binding affinity (–8.4 kcal/mol), lowers the ATP level with a minimal concentration as compared to 64-9C. Hence, we picked 64-9D for the evaluation of other microbial activities. The effectiveness of the hit molecule 64-9D was confirmed through the inhibitory assay against *Mtb* mc² 6230. Importantly, the bioenergetic machinery of this strain is identical to the virulent strain *Mtb* H37Rv. Encouraged by these initial findings, we have synthesised a series of structural analogues of 64-9D. We have synthesised a focused library of 8-position ester-substituted quinoline derivatives PPQN-2FQN. All the synthesised quinoline-based ester derivatives (PPQN-2FQN) were evaluated for their antimycobacterial activity using the liquid broth microdilution assay against the *Mtb* mc² 6230 strain. The MIC values are listed in (Figure 2C) and the structure–activity relationship (SAR) was deduced accordingly.

### Synthesis

Based on the ATP-depleting activity and inhibitory effects against *Mtb*, a series of structural analogues of 64-9D were synthesized to further improve therapeutic potential. A series (PPQN-2FQN) of ten quinoline-based esters was synthesised through a one-step esterification method (Figure 1). In this approach, 8-hydroxyquinoline was reacted with a diverse set of carboxylic acids, including aliphatic, cyclic, and substituted aromatic acids, employing 1-ethyl-3-(3-dimethylaminopropyl) carbodiimide hydrochloride (EDC⋅HCl) as the coupling agent. The reaction was facilitated by the addition of 1-hydroxybenzotriazole hydrate (HOBt), catalytic 4-dimethylaminopyridine (DMAP) and triethylamine (TEA) under ambient conditions. This method afforded the corresponding esters in moderate to good yields.(Pieters *et al*, 2018; Islam *et al*, 2023)

**Figure 1:**
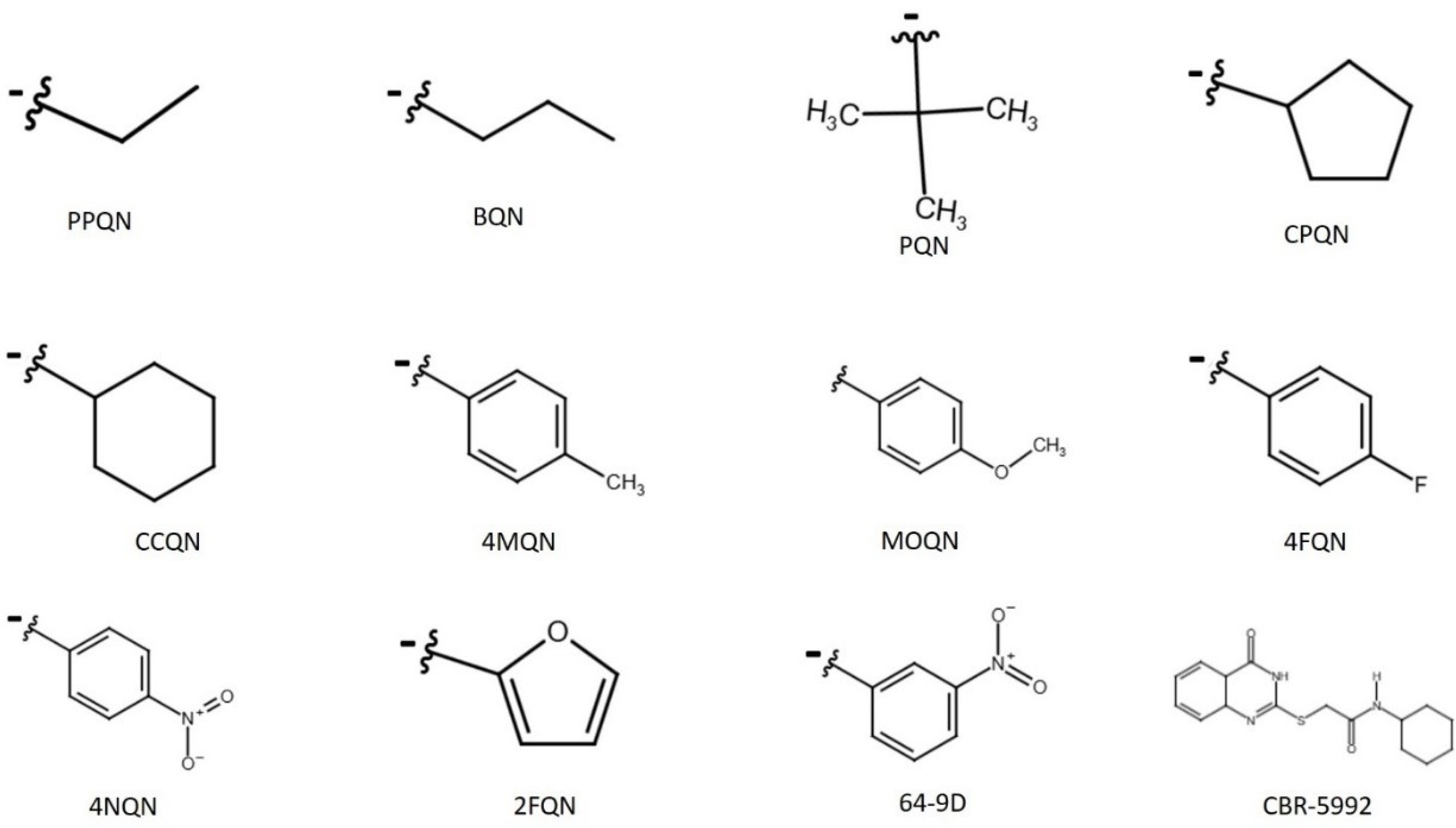
Synthesis for optimization: Chemical structures of synthesized analogues of 64-9D highlighting site-specific modifications based on SAR analysis.

**Figure 2:**
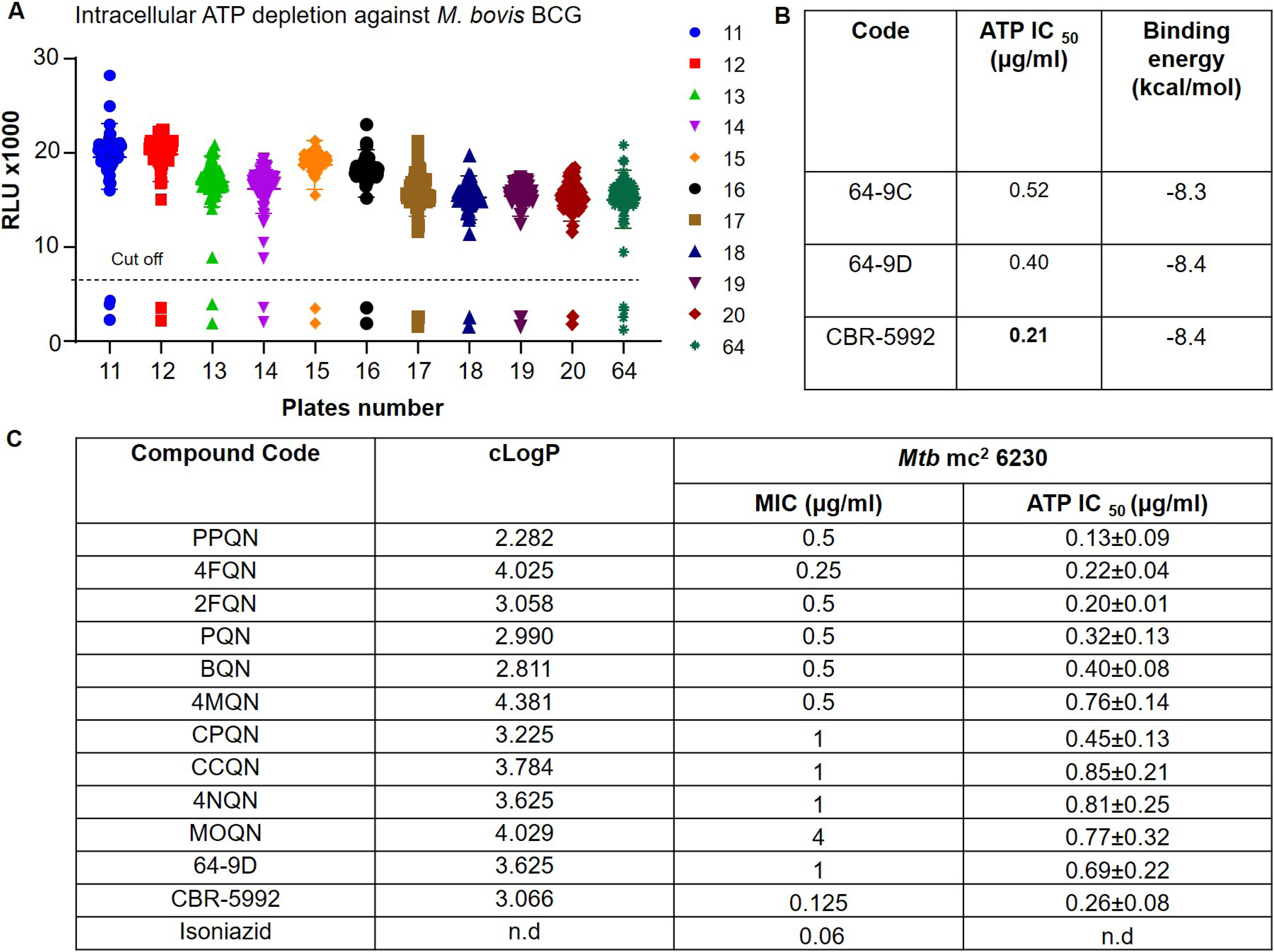
Anti-tubercular activity-based evaluation for lead optimization: ATP depletion assay for screening of molecules against mycobacteria using luciferase-based assay (A). Determination of ATP IC_50_ for most active scaffolds. ATP levels were measured in *Mtb* mc² 6230 using a luciferase-based assay (B). MIC and ATPIC50 of structural analogues of 64-9D against auxotrophic stains of *Mtb* H37Rv (C). The data were expressed as mean ± SDs (n=3), with triplicate measurements for each compound concentration. The experiment in (A) was conducted as a single replicate. The SD of the three biological experiments from a single experiment is shown by the error bar.

**Scheme 1.**
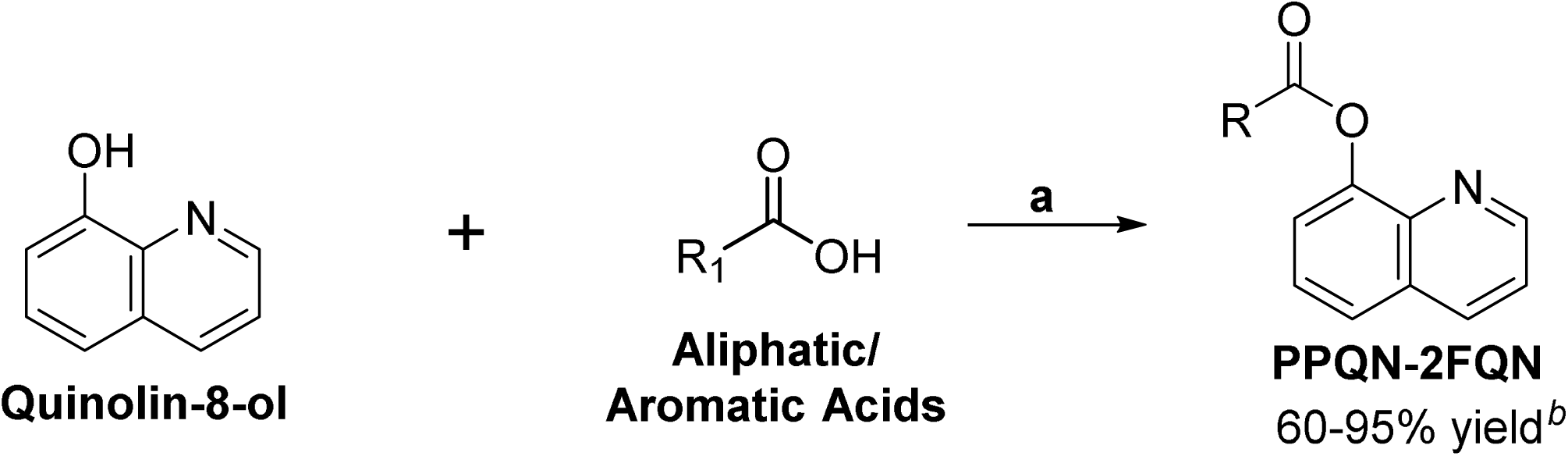
Synthetic approach for (Quinolin-8-yl) ester derivatives **PPQN-2FQN**. **^a^Reagents and conditions**: (a) 8-hydroxyquinoline (1 mmol), carboxylic acids (1 mmol), EDC.HCl (1.5 mmol), HOBt (1.5 mmol), DMAP (0.1mmol) and TEA (1 mmol), DCM (5 mL, for 1 mmol reaction), room temperature, ^b^isolated yield.

Among these compounds, the aliphatic-substituted quinoline-8-yl ester derivatives possessed linear or branched aliphatic side chains, like propionate (PPQN), butyrate (BQN), and pivalate (PQN), exhibited notable potency with MIC values of 0.5 µg/mL. This suggests that lipophilic aliphatic groups, irrespective of their branching pattern, are well tolerated and favourable for antimycobacterial activity, likely due to enhanced membrane penetrartion or target binding affinity. Also, the cycloalkyl-substituted ester derivatives of quinoline-8-yl, like cyclopentyl carboxylate (CPQN) and cyclohexyl carboxylate (CCQN) showed identical MIC values of 1 µg/mL. Next, phenyl ester derivatives, with elctron withdrawing groups like p-fluoro (4FQN) and p-nitro (4NQN) displayed the MIC of 0.25 µg/mL and 1 µg/mL, respectively. Phenyl ester derivativatives with electron donating group like p-methyl (4MQN) and p-methoxy (4-MOQN) displayed MIC of 1 µg/mL and 4 µg/mL, respectively. At last, heteroaryl ester derivative 2FQN features a furan-2-yl side chain exhibits an MIC of 0.5 µg/mL.

After evaluating the activity of all synthesised compounds against the *Mtb* mc² 6230 strain and other resistant strains, the most active compound was found to be 4FQN, followed by 2FQN. Based on their superior activity profiles, these two molecules were selected for further detailed studies. CBR-5992, a known NDH-2 inhibitor, was utilized as a control to assess ATP depletion. Meanwhile, isoniazid, rifampicin, and CBR-5992 were employed as standard drugs for determining the minimum inhibitory concentration (MIC), with DMSO serving as a drug-free control.

### Quinolone lead molecules are equipotent against multidrug resistant tuberculosis

Though NDH-2 is an integral part of the ETC and ATP is the end product of the Ox-Phos pathway, we first evaluated the ATP depletion property of the synthesized molecules. The intracellular ATP depletion assay was conducted in a 96-well white flat-bottom plate (Corning) to assess the effectiveness of the test actives against susceptible *Mtb* mc² 6230,(Kalia *et al*, 2017b) revealing that all molecules exhibit comparable efficacy in reducing intracellular ATP levels. This suggests their equivalent potency in disrupting cellular energy metabolism in *Mtb* (Figure 2C). Hence it was very interesting to evaluate the other antibacterial activity of these compounds. The active molecules were evaluated for their growth inhibition ability against multi-drug-resistant strains of *M. tuberculosis* viz. *Mtb* mc² 8243, *Mtb* mc² 8247, and *Mtb* mc² 8259 strains. We observed that the molecules were inhibiting the bacterial growth against different strains of the *Mtb* at a good range (0.25-4 µg/ml) of concentration (Figure SI 1A). Additionally, these test actives were also able to inhibit the bacterial growth and deplete the whole cell intracellular ATP against *Mtb* H37Ra (Figure SI 1A). Interestingly, among the synthesized analogues, 06 compounds (namely PPQN, 4FQN, 2FQN, PQN, BQN, 4MQN) deplete the intracellular ATP abruptly and equipotent against mono and multi-drug-resistant strains of *Mtb*.

## Target Validation: *In-vitro* & *In-silico* approach

### Quantitative assessment of NADH: NAD⁺ ratio using Peredox mCherry Biosensor

NADH dehydrogenase (NDH-2) is the first enzyme in the electron transport chain (ETC) of the oxidative phosphorylation pathway, facilitating the transfer of electrons through the oxidation of NADH to NAD⁺. In *Mtb*, uninterrupted ATP synthesis is critically dependent on functional NDH-2 activity.(Beites *et al*, 2019) To evaluate the target specificity of the active analogues of 64-9D toward NDH-2, a strain of *Mtb* mc^2^ 6230 transformed with Peredox-mCherry biosensor was used.(Bhat *et al*, 2016) The biosensor containing strain of *Mtb* was exposed to desired concentrations of test molecules and on binding to NDH-2, the enzymatic conversion of NADH to NAD⁺ was disrupted that resulted in intracellular accumulation of free NADH. This excess NADH interacted with the circularly permuted T-sapphire domain of the reporter system of biosensor, leading to an increase in green fluorescence intensity, that served as a quantitative proxy for NDH-2 inhibition (Figure 3A (i-vi)). The change in Green/Red ratio in test samples was compared with untreated control. It was observed that the effect of all test molecules on Green/Red ratio was in concentration dependent manner and at 4×MIC values the changes was similar or close to the values obtained for CBR-5992 at its 5×MIC, which is a known NDH-2 inhibitor.(Bhat *et al*, 2016; Boshoff *et al*, 2008; GOPINATHAN *et al*, 1963) There was no significant effect of CLFZ or rotenone (NDH-1 inhibitor) on Green/Red ratio as both do not directly impact the functioning of NDH-2.

**Figure 3:**
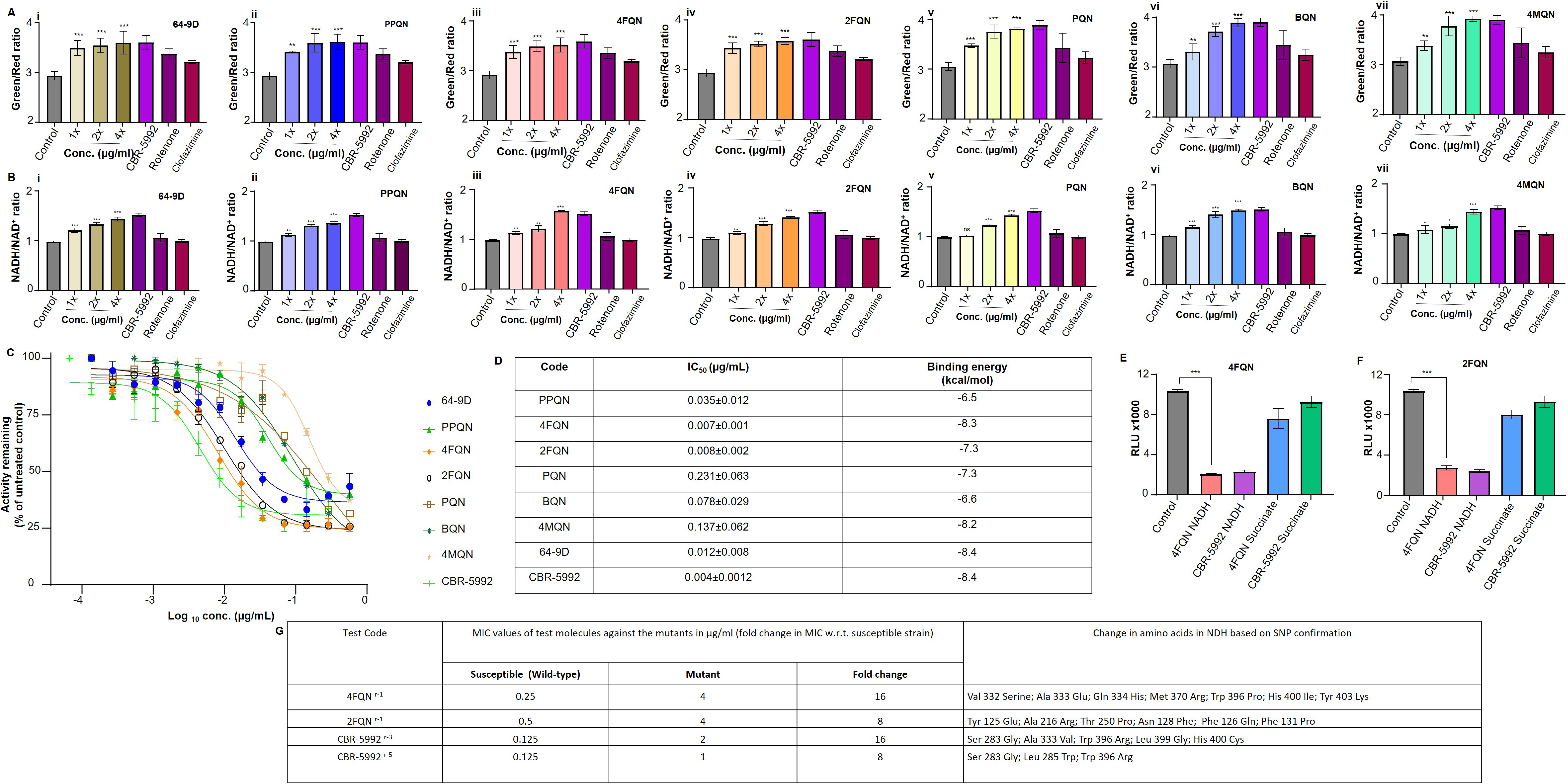
Validation of Type-II NADH as a biomolecular target of the shortlisted molecules: The pMV762-Peredox-mCherry expressing *Mtb* mc² 6230 strain is sensitive to variations in the NADH: NAD^+^ ratio. The molecules exhibit a dose-dependent inhibition of the NADH dehydrogenase complex (A(i-vi)). Reconfirmation and evaluation of disruption in NADH: NAD^+^ ratio using an alternative fluorescence-based assay (B(i-vi)). Enzyme inhibition assay using purified recombinant *Mtb*-NDH protein and determination of the IC_50_ (C, D). *In-silico* interactions studies using NDH protein as a ligand for determination of binding energy of the molecules (D). 4FQN and 2FQN significantly suppress ATP production compared to the untreated control when NADH serves as the electron donor (E) and (F) respectively. However, this inhibition is not observed when succinate is used as the electron donor in inverted membrane vesicles (IMVs). SNPs identification in *ndh* gene for one step mutants raised against 4FQN, 2FQN, and CBR-5992 (G). The data were expressed as mean ± SDs (n=3), with triplicate measurements for each concentration. Statistical significance was assessed using one-way ANOVA followed by Dunnett’s test, where \**P* < 0.05, ** *p* < 0.01, and *** *p* < 0.001 compared to the untreated control

### Quantification of change in Redox Homeostasis based on NADH/NAD⁺ ratio using a fluorescent probe

The redox homeostasis in *Mtb* is essential for its survival under various environmental conditions inside the host. There are multiple redox couples that work independently or linked to others. The NADH/NAD^+^ coenzyme system is required for catabolism and NAD^+^ is an efficient electron sink which is used by the bacteria for as a cofactor in several oxidizing reactions.(Xu *et al*, 2023b) A constant level of NADH is essential for mycobacterial growth, whereas the concentration of NAD^+^ is variable and any change in the ratio will affect the growth of bacilli. The typical ratio of NADH/NAD^+^ required for normal growth ∼1:3 to 1:10, but when bacteria is shifting from aerobic to anaerobic conditions a higher ratio NADH/NAD^+^ is generated and this is because of depletion of the NAD^+^ pool, and that is maintained by type II NADH dehydrogenase. Finally, the redox homeostasis plays an important role in ATP generation which is essential to perform all cellular activities. Hence, distributing of the function of NDH-2 will severely impair the energy production, and making it a strategic target for therapeutic intervention (Figure 3B (i-vi)). Therefore, the ratio of NADH/NAD^+^ was quantified for *Mtb* after exposure to probable inhibitors of NDH-2 using a fluorescence-based assay system. The effect of test molecules was quantified and NADH accumulation was observed in dose-dependent manner, suggesting effective inhibition of NDH-2 enzymatic activity. These findings supported our hypothesis and further validated NDH-2 as a viable candidate for therapeutic intervention in *Mtb*.

### Interaction studies using *in-silico* tools to identify binding sites and affinity with target protein

In the present study, following the synthesis and *in-vitro* evaluation of quinoline-based ester compounds for their anti-tubercular activity, molecular docking was conducted to investigate the potential binding interactions of the structural analogs including 64-9D. The Known inhibitor of NDH-2 (CBR-5992) served as positive control. The crystal structure of *Mycobacterium smegmatis* NDH-2 (PDB ID: 9MQZ) was used a target protein(Liang *et al*, 2025) and to perform docking simulations were using AutoDock Vina version 1.1.2,(Trott & Olson, 2010) focusing on the predefined binding site of the 2-mercapto-quinazolinone inhibitor (Figure SI 2).

*In-silico* molecular docking studies revealed that the test compounds exhibited favourable binding affinities with the target protein, suggesting a strong interaction potential with the ligand. All the molecules demonstrated binding energies within a therapeutically relevant range. These findings support their candidacy for further experimental validation and optimization (Figure SI 2).

### Target Validation using recombinant purified NDH protein

The pYUB-C-term-*Mtb-ndh* construct was electroporated into *M. smegmatis* mc² 4517, and positive clones were selected on hygromycin B (50 µg/mL). Protein was extracted via detergent solubilization and purified by affinity chromatography using imidazole gradients (20 mM wash, 150 mM elution), with SDS-PAGE confirming elution efficiency (Figure SI 3A).(Lu *et al*, 2022; Dam *et al*, 2022)

### Inhibition of recombinant His-*Mtb*-NDH by selected molecules

The enzyme inhibition assay was performed against the recombinant purified enzyme which will confirm NDH-2 as a target protein for the test molecules. The enzyme inhibitory profiling revealed that compounds 4FQN and 2FQN exhibited IC₅₀ values of 0.008 µg/ml and 0.007 µg/ml respectively. The enzyme assay results confirmed 4FQN as the most effective inhibitor of NDH-2 in terms of target engagement followed by 2FQN (Figure 3 & D).

To further characterize the inhibition mechanism of the most promising 64-9D analogs i.e. 4FQN and 2FQN, enzyme kinetics studies were conducted using double-reciprocal (Lineweaver–Burk) plots. These analyses allowed to evaluate the type of inhibition modes and estimation of kinetic constants, providing deeper insight into compound–target interactions. The representative plots for 4FQN, 2FQN, and CBR-5992 were generated based on substrate velocity data in the presence of varying inhibitor concentrations and the data obtained revealed uncompetitive type inhibition by proposed NDH-2 inhibitors for *Mtb* (Figure SI 3B,C,D,&E).(Yano *et al*, 2006; Murugesan *et al*, 2018)

### NADH-fueled electron transport in IMVs: A tool to pinpoint inhibition targets

Given the central role of NDH-2 in the ETC and its contribution to ATP synthesis via oxidative phosphorylation, the ATP-depleting potential of the test compounds was evaluated using inverted membrane vesicles (IMVs). To identify the electron entry point, ATP synthesis was measured in the presence of two distinct electron donors: NADH and Succinate. The results demonstrated that when NADH was supplied as the electron donor, the test compounds significantly inhibited ATP synthesis compared to the drug-free control. In contrast, no substantial inhibition was observed with succinate, indicating that the compounds specifically target the NADH-dependent entry pathway, consistent with NDH-2 inhibition. Known NDH-2 inhibitor i.e CBR-5992 was included as positive control to validate the assay (Figure 3E & F).(Murugesan *et al*, 2018)

### Susceptibilities of mutants, cross-resistance to test drugs and mutations in NDH-2 encoding gene

One-step mutants against most active structural analogues of 64-9D i.e 2FQN and 4QN including CBR-5992, a known inhibitor of NDH-2 were raised, evaluated for susceptibility and those showed more than 8 fold change in MIC values against respective test drugs were shortlisted and sequenced for identification of SNPs in *ndh* gene.(Saha *et al*, 2024c) Sequencing of resistant clones revealed point mutations at multiple loci within the *ndh* gene, supporting the hypothesis that CBR-5992 and the test compounds interact with distinct binding sites on NDH-2. Further, the characterized mutants for one test drugs were tested for susceptibility against all shortlisted test drugs to confirm cross-resistance if any (Figure 3G). No significant alterations in MIC values were detected for the test compounds against reciprocal mutants, indicating the absence of cross-resistance. Furthermore, the test drugs maintained their efficacy against CBR-5992 mutants.

### Bactericidal potency of 4FQN and 2FQN against replicating and non-replicating *Mtb*

Anti-mycobacterial agents can be either bacteriostatic or bactericidal. The bacteriostatic agents may arrest the bacterial growth, but for complete eradication of bacterium from the site of infection bactericidal response is needed. The mid log phase actively replicating *Mtb* mc^2^ 6230 when exposed to the shortlisted test molecules (4FQN and 2FQN) at 1-4×MIC concentrations for 5 and 10 days. both the compounds i.e 4FQN and 2FQN exhibits time and concentration dependent killing w.r.t. initial inoculum.(Harbut *et al*, 2018; Awasthy *et al*, 2014; Fisher *et al*, 2009; Heikal *et al*, 2014; Xu *et al*, 2023a) The bacterial load for both the test molecules was reduced by 1.5 Log_10_ CFU/ml (MBC_90_) at 4×MIC values after 5 days of exposure and same effect was observed at 2×MIC after 10 days of exposure (Figure 4A). Also, ∼ 3Log_10_ CFU/ml (∼99.9%) reduction in bacterial load was observed at 4×MIC after 10 days of exposure revealing that test were more effective at 4× MIC. The CBR-5992 (1.25 µg/mL), a well-characterized NDH-2 inhibitor, was included as a positive control.

**Figure 4:**
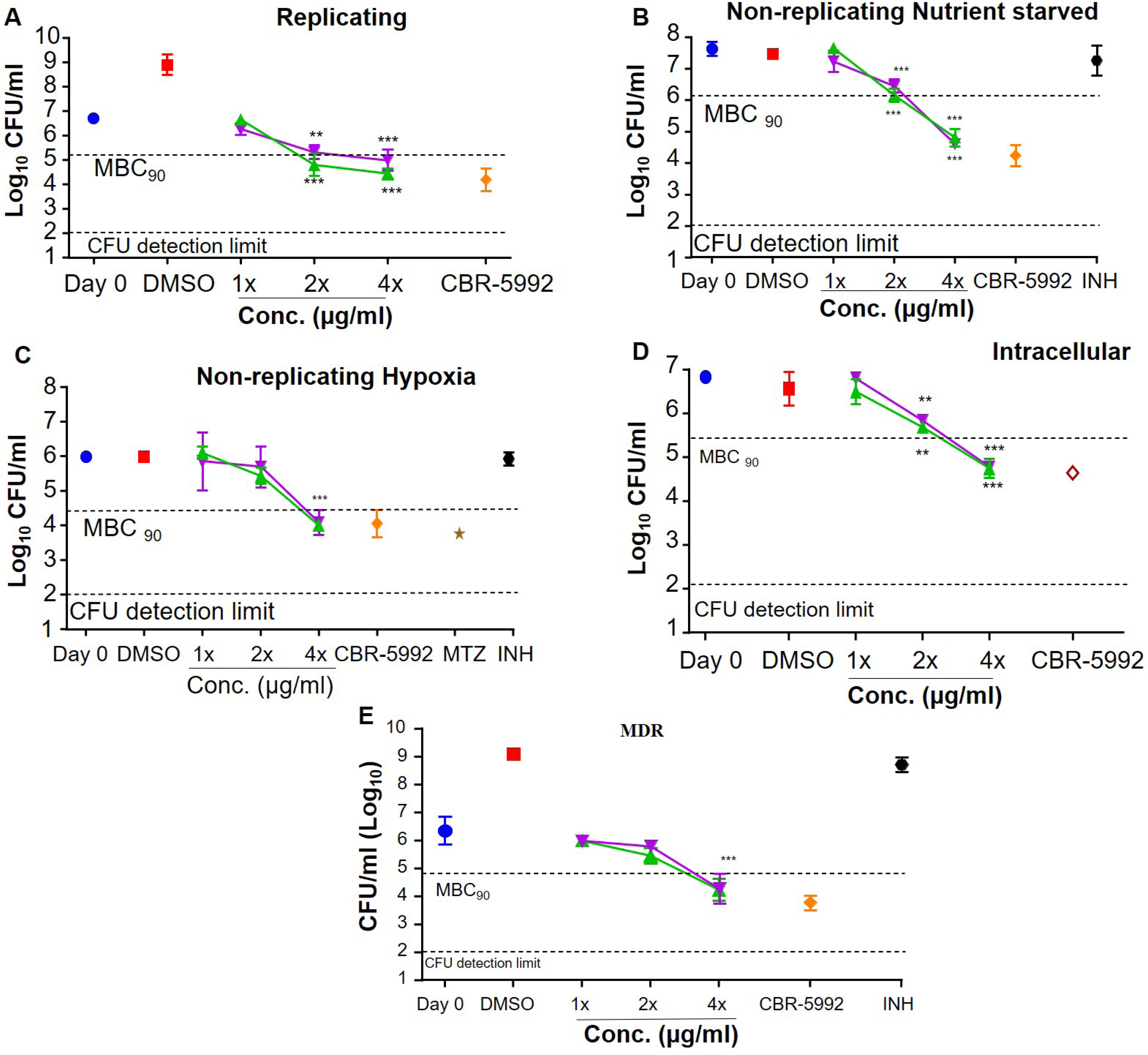
The type II NADH inhibitors were bactericidal against replicating, non-replicating, and resistant Mycobacteria: The bactericidal activity of 4FQN and 2FQN (1x-4x MIC) against replicating (A), non-replicating nutrient starved (B), non-replicating-hypoxia (c), intracellularly-invaded mycobacteria, and multidrug resistant strain of mycobacteria. CBR-5992, INH and MTZ served as conditional drug controls. On Day 0, the initial inoculum was established, while DMSO was used as a drug-free control. Bacterial viability was determined by counting colony-forming units (CFU) on 7H10-agar plates following a 10-day incubation period. Green triangles denote 4FQN, while purple triangles represent 2FQN. The initial inoculum at day 0 is shown as blue circles, DMSO as red squares, CBR-5992 as orange diamonds, INH as black circles, and MTZ as brown stars. The dotted line represents a 90% reduction in CFU/ml compared to the initial inoculum at Day 0. Data were expressed as mean ± SDs (n=3), with triplicate measurements for each compound concentration. Statistical significance was assessed using one-way ANOVA followed by Dunnett’s test, where *** *p* < 0.001 vs. Day 0 initial inoculumThe SD of the three biological experiments from a single experiment is shown by the error bar.

### Killing efficacy of test molecules against non-replicating nutrient starved and hypoxic form of mycobacteria

Test actives were evaluated for their killing efficacy against non-replicating nutrient starved and hypoxic *Mtb* mc^2^6230. The non replicating bacilli were exposed to different concentrations of the test molecules in multiples of their MICs. Isoniazid served as a negative drug control for nutrient starved nonreplicating mycobacteria, metronidazole as a positive control for hypoxic conditions, and CBR-5992 as a known NDH-2 inhibitor for both the conditions.(Kalia *et al*, 2017b; Lee *et al*, 2021; Kalia *et al*, 2023) The results depicted that both test molecules at 4×MIC decreased the bacterial load by more than 1.5 Log_10_ CFU/ml (Figure 4B & C) and confirmed their efficacy under non-replicating conditions as well, proving them as a better candidate for anti-tb drug development because they target heterogenous population of *Mtb*.

### Time-kill kinetic of NDH-2 inhibitors against replicating mycobacteria

Taking lead from MBC experiments, the same molecules at concentrations in multiples of MIC ranging from 1 4×MIC were tested for evaluation of time and concentration dependent killing efficacy against *Mtb* mc^2^6230 in a time kill assay. The CFUs were determined at regular intervals 03 days until day 15. Both the 4FQN and 2FQN exerted a strong and rapid bactericidal activity that led to a reduction of 3 Log_10_CFU/ml within 9 days at 4×MIC concentration (Figure SI 4A & B). The test molecules managed to reduce the bacterial load by 3 Log_10_ CFU/ml even at 2×MIC concentration after 12 days of exposure. The results obtained confirmed the time and concentration dependent killing of bacilli by test molecules.(Shetye *et al*, 2020; Saha *et al*, 2024c)

### Intracellular killing potential of NDH-2 inhibitors

The intracellular activity of the compounds was done to evaluate their ability to penetrate the macrophage and kill the bacteria. We further assessed the MICs of the NDH-2 inhibitors against *Mtb* H37Ra and observed values comparable to those obtained with *Mtb* mc² 6230. The *Mtb* H37Ra infected monolayers of THP-1 cell lines were treated for 5 consecutive days with multiple concentrations of the test molecules. The results obtained were analysed using Log_10_ CFU/ml, compared with initial inoculum, untreated and control drugs (Figure 4D). It was observed that at 4×MIC both 4FQN and 2FQN significantly reduced the bacterial load by ∼2Log_10_ CFU/ml and confirmed that mammalian cells are permeable to these molecules, which is one of the necessary properties of drug like candidates.

### *In-vitro* intracellular ATP depletion assay against nutrient starved *Mtb* mc^2^ 6230

Next, the effect of 4FQN and 2FQN treatment on ATP homeostasis was measure using nutrient starved *Mtb* mc^2^ 6230.(Berube *et al*, 2019) ATP depletion assay was executed in a 96 well white flat bottom plate (Corning USA) to evaluate the efficacy of the test actives to deplete the intracellular ATP against nutrient starved *Mtb* mc^2^ 6230.(Kalia *et al*, 2017b) Previously the test actives achieved the cidality against the different phenotypic conditions and here also they were able to inhibit the whole cell intracellular ATP synthesis within a good range of concentrations (Figure SI 4C &D).

### Killing Efficacy against multidrug resistant stains

Anti-mycobacterial agents can be either bacteriostatic or bactericidal. The bacteriostatic agents may arrest the bacterial growth, but for complete eradication of bacterium from the site of infection bactericidal response is needed. The mid log phase actively replicating *Mtb* mc^2^ 8259 when exposed to the shortlisted test molecules at concentrations in multiples of MIC values ranging from 1×MIC to 4×MIC for 10 days (Figure 4E). It was observed that the killing effect of both the shortlisted test molecules i.e 4FQN and 2FQN was time and concentration dependent w.r.t. initial inoculum.(Harbut *et al*, 2018; Awasthy *et al*, 2014; Fisher *et al*, 2009; Heikal *et al*, 2014; Xu *et al*, 2023a) The bacterial load for both the test molecules was reduced by 1.5 Log_10_ CFU/ml at 4× MIC values after 10 days of exposure revealing that test molecules are more effective at 4× MIC and like other ETC inhibitors, NDH2 inhibitors also need more time to kill the bacteria. The CBR-5992 (1.25 µg/mL), a well-characterized NDH-2 inhibitor, and isoniazid was included as a positive control.

### Oxygen consumption inhibition using methylene blue as an oxygen sensor

The test compounds were assessed for their ability to inhibit oxygen consumption, with methylene blue serving as a qualitative indicator of the dissolved oxygen levels in the culture media.(Moosa *et al*, 2017; Xu *et al*, 2023b) The compounds effectively inhibit oxygen consumption, maintaining a blue colour, in contrast to other drugs that do not completely block the electron transport chain (ETC) and turn colourless. Using methylene blue as an oxygen probe, we observed that over 96 hours, BDQ therapy significantly reduced oxygen utilization, whereas Q203 treatment showed no effect. These findings from the methylene blue assay confirm that our test actives were capable of inhibiting the oxygen consumption process in the ETC of *Mtb* (Figure 5A). As there was no alternative pathway to share the electrons to the ETC, targeting the NDH-2 inhibition, completely inhibits the respiration process of *Mtb.* Which additionally support our hypothesis that these compounds inhibit the oxidative phosphorylation pathway of *Mtb.* Previous studies reported that those molecules were inhibiting the respiration process completely; they remain as blue.(Kalia *et al*, 2017a; Lee *et al*, 2019)

**Figure 5:**
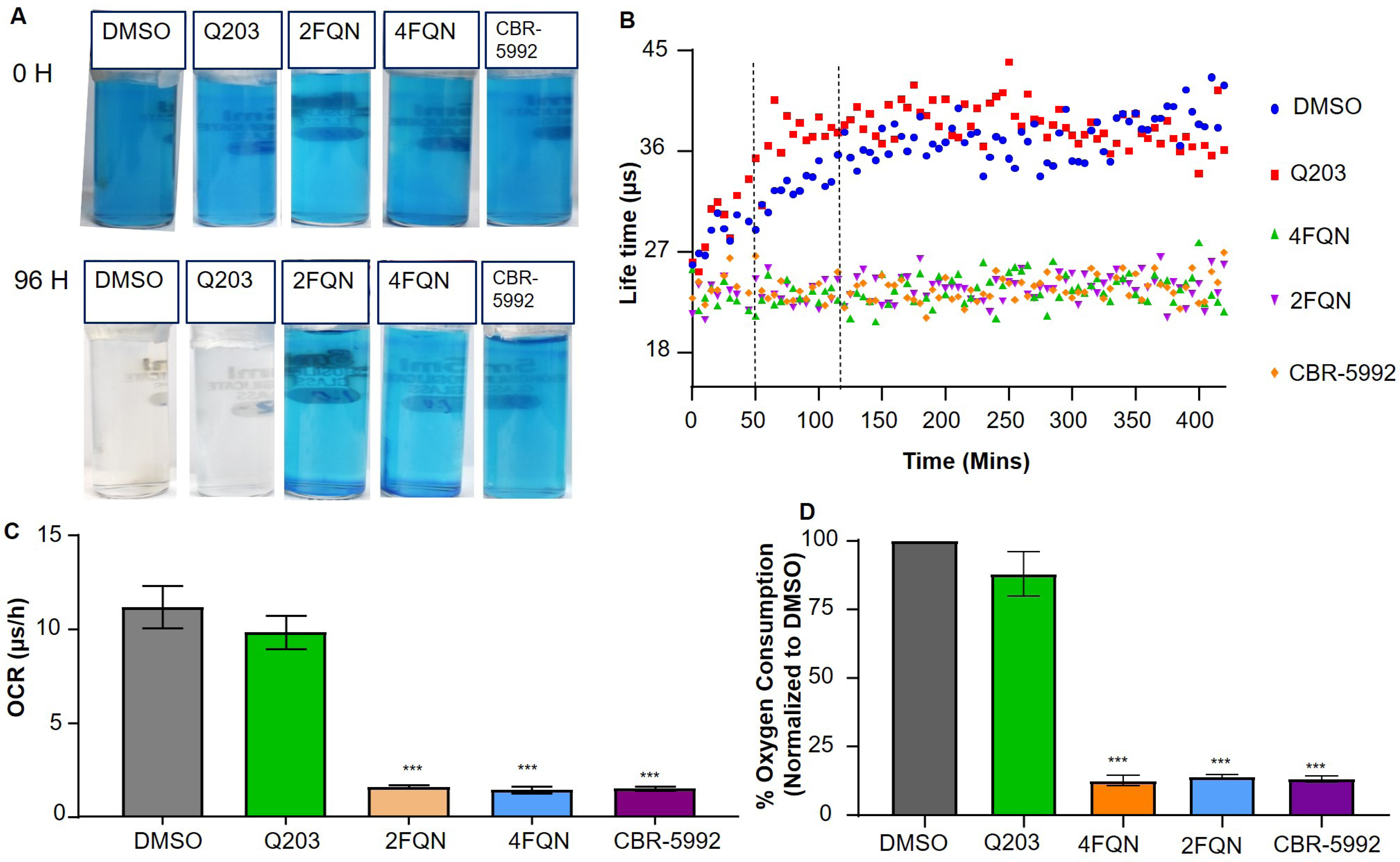
The inhibitors of type II NADH dehydrogenase inhibits the respiration in replicating mycobacteria: The oxygen consumption assay in *Mtb* mc² 6230 using methylene blue oxygen sensor at 0.001% (A). MitoXpress, an alternative fluorescence-based assay was used for kinetics of oxygen consumption and the fluorescence (Ex380, Em650) was recorded over a 420-min period. The bacteria were incubated with DMSO (blue circles), Q203 treated (red square), 4FQN (green triangles), 2FQN (purple triangles), CBR-5992 (orange diamonds) and fluorescence was recorded. The results were represented in form life time (B), oxygen consumption rate (C), and % O_2_ consumed w.r.t untreated control i.e. DMSO (D). The experiments were performed in triplicate and repeated at least once. Data are expressed as the mean ± SDs of triplicates foreach concentration of a representative experiment. Statistical significance was assessed using one-way ANOVA followed by Dunnett’s test, where \**p* < 0.05, ** *p* < 0.01, and *** *p* < 0.001 compared to the DMSO control. The SD of the three biological experiments from a single experiment is shown by the error bar.

### Oxygen consumption assays using the MitoXpress® oxygen probe

The test compounds were assessed for their ability to inhibit oxygen consumption using the MitoXpress probe to measure dissolved oxygen levels in the culture media of whole cells over a short period.(Roy *et al*, 2026) Under the experimental setup, Q203 showed no significant effect on oxygen respiration in the replicating strain, whereas BDQ caused complete inhibition of oxygen consumption. Notably, the test molecules significantly suppressed dissolved oxygen consumption compared to the drug-free control. In this assay, CBR-5992 and Q203 were used as drug controls (Figure 5B, C & D). *Mtb* mc² 6230 cells were incubated with the Mito-Xpress oxygen probe in the presence of 1% DMSO, 100 nM Q203, and test compounds at 10×MIC concentration along with CBR-5992. The kinetics of oxygen consumption was monitored by measuring fluorescence (Ex380, Em650) over a 420-minute period. Kalia et al. reported similar findings when evaluating molecules that completely inhibit respiration.(Kalia *et al*, 2017a)

### *In-vitro* checkerboard assay with Q203 and bedaquiline against replicating *Mtb* mc^2^ 6230

To assess the potential synergistic effects of novel quinoline derivatives as NDH-2 inhibitors in combination with established antimycobacterial agents, we conducted *in-vitro* checkerboard assays using 4FQN and 2FQN against the susceptible *Mtb* mc² 6230 strain. Two distinct respiratory inhibitors Q203 and bedaquiline (BDQ) were selected for combination studies based on their complementary mechanisms of action. Q203 targets the Cytochrome *bc1:aa3* super complex, a critical component of the mycobacterial electron transport chain. When paired with Q203 (Figure SI 5 (i&iii)) or BDQ (Figure SI 5 (ii&iv)) both NDH-2 inhibitors demonstrated additive effects.

This suggests that simultaneous inhibition of multiple respiratory nodes may enhance metabolic disruption and contribute to improved bactericidal activity. The additive interaction supports the rationale for combinational therapy aimed at boosting lesion sterilization and minimizing the emergence of resistance. BDQ inhibits the ATP synthase complex, thereby impairing energy generation in *Mtb*. Its combination with 4FQN and 2FQN also yielded additive effects, with enhanced inhibitory activity observed across the tested concentrations. These results further reinforce the therapeutic potential of targeting distinct metabolic pathways in tandem, offering a complementary strategy to augment treatment efficacy.

### Extended growth inhibition of *Mtb*: post drug withdrawal

The post-antibiotic assay is performed to evaluate the duration of bacterial growth suppression after brief antibiotic exposure. It helps determine the post-antibiotic effect (PAE), providing insights into dosing intervals and therapeutic efficacy. The post-antibiotic effect of the tested molecules was assessed using *Mtb* mc^2^ 6230 to determine the duration of their efficacy following drug removal.(Chan *et al*, 2001, 2004; Shetye *et al*, 2020) The experiment revealed that at a 4×MIC concentration, compound 4FQN exhibited prolonged activity for up to 96 hours, while 2FQN extended its effect for 48 hours. CBR-5992, used as a standard, demonstrated a similar extension of activity for 48 hours at the same concentration (Figure SI 1B).

### Frequency of mutation generation

The concentration of test drug at which there was no colony after incubation was considered as its MPC.(Kalia *et al*, 2012) The mutation prevention concentration (MPC) assay revealed distinct potencies between the tested compounds. While 4FQN inhibited *Mtb* mc² 6230 growth at 16× MIC, 2FQN required 32× MIC to achieve complete suppression. This suggested that 4FQN may have a higher barrier to resistance development compared to 2FQN. These findings underscored the importance of evaluating MPC alongside MIC for selecting compounds with favorable resistance suppression profiles (Figure SI 1C).

### Safety assessment using HepG2 cell lines

Cytotoxic effect of the compounds was tested on HepG2 cell line. Percentage cell viability of HepG2 cell line was estimated by using Trypan blue dye exclusion technique. Viable cells were able to exclude the dye and therefore did not take up blue color whereas dead cells were stained blue. These NDH-2 inhibitors have a good range of cytotoxicity profile (Safety index >10) and safe to use (Saha *et al*, 2025) (Figure SI 1D).

### Molecular Docking Study

In the present study, following the synthesis and in vitro evaluation of quinoline-based ester compounds for their anti-tubercular activity, molecular docking was conducted to investigate the potential binding interactions of the most active derivatives (4FQN and 2FQN), 64-9D (parent compound) and the standard compound CBR-5992. For this purpose, the crystal structure of *Mycobacterium smegmatis* NDH-2 (PDB ID: 9MQZ) was utilised.(Liang *et al*, 2025) The docking simulations were carried out using AutoDock Vina version 1.1.2,(Trott & Olson, 2010) focusing on the predefined binding site of the 2-mercapto-quinazolinone inhibitor. The key binding interactions are depicted in (Figure SI 2). while the amino acid residues involved in ligand binding are summarised.

The binding interactions, providing insight into the probable types of interactions between the amino acid residues at the active site and the structural motifs of the synthesised derivatives, thereby highlighting the essential features required for effective binding. All four ligands consistently formed conventional hydrogen bonds with THR^A:178^ and FAD^A:501^. Specifically, 4FQN, 2FQN, and 64-9D each formed strong hydrogen bonds with THR^A:178^, indicating the importance of this residue in stabilising the ligand. In the case of CBR-5992, it forms hydrogen bonds with GLY^A:276^ and GLY^A:328^, offering an extended polar interaction network due to its elongated scaffold (Figure SI 2).

In terms of π-π stacking interactions, FAD^A:501^ seems central across all ligands. 4FQN, 2FQN, and 64_9D displayed π-π stacked or T-shaped interactions with FAD^A:501^, signifying efficient aromatic alignment with the flavin ring system. Although CBR-5992 lacked classical π-π stacking, it participated in an amide-π stacked interaction, reflecting a unique binding motif. Hydrophobic interactions, such as π-π-alkyl and alkyl interactions, were also observed. PRO^A:177^ and LYS^A:136^ are repeatedly involved in π-π-alkyl or alkyl interactions in 2FQN, 64-9D and CBR-5992, providing hydrophobic stabilisation. VAL^A:329^ was involved in alkyl or π-alkyl interaction in both 64-9D and CBR-5992.

4FQN, the most active compound in the series, uniquely featured an additional hydrogen bond with MET^A:135^ through the fluorine atom of the ring, absent in the others, which may explain its superior binding affinity. It also combined all key stabilising interactions observed in other compounds of the series. In contrast, the parent compound 64-9D maintained a basic but stable interaction pattern, lacking the extended hydrogen bond network seen in 4FQN. Finally, CBR-

5992, despite its bulkier structure, showed extensive polar and hydrophobic contacts across multiple residues (e.g., GLY^A:119^, GLY^A:276^, VAL^A:329^, FAD^A:501^), supporting its stability as a standard, though lacking key interactions like THR^A:178^ binding seen in the most active compounds.

### *In-Silico* physicochemical properties and ADME profiling

The pharmacokinetic and drug-likeness characteristics of 4FQN and 2FQN were predicted using SwissADME and pkCSM platforms(Daina *et al*, 2017; Pires *et al*, 2015). Both molecules exhibited excellent human intestinal absorption, with predicted absorption values of 96.95 % (4FQN) and 98.39 % (2FQN), indicating high oral bioavailability potential. Neither compound was identified as a P-glycoprotein substrate, suggesting minimal risk of efflux-mediated reduction in intracellular concentration. In terms of distribution, the predicted volume of distribution (log VDss) values (4FQN: –0.125; 2FQN: 0.11) were within the favourable range for adequate tissue penetration. Both compounds demonstrated the ability to cross the blood–brain barrier (log BB: 0.594 and 0.224, respectively) and displayed acceptable CNS permeability (log PS: –1.431 and –2.840), supporting potential central nervous system activity. Metabolic stability predictions indicated that both compounds are substrates for CYP3A4 but not for CYP2D6, which may facilitate predictable metabolic pathways. Clearance values (log mL/min/kg) of 0.55 for 4FQN and 0.796 for 2FQN suggest efficient elimination via hepatic and renal routes. From a physicochemical perspective, both derivatives showed hydrogen bond donor (0) and acceptor (4) and topological polar surface area (TPSA) values within the optimal range for drug absorption and permeability (4FQN: 39.19 Å²; 2FQN: 52.33 Å²). Both compounds fully follow Lipinski’s Rule of Five and exhibited zero PAINS alerts. Overall, the ADME and physicochemical profile (Table 1) of 4FQN and 2FQN highlights their promising oral absorption, favourable distribution, CNS penetration potential, and balanced physicochemical properties, making them attractive candidates for further optimisation.

**Table 1:**
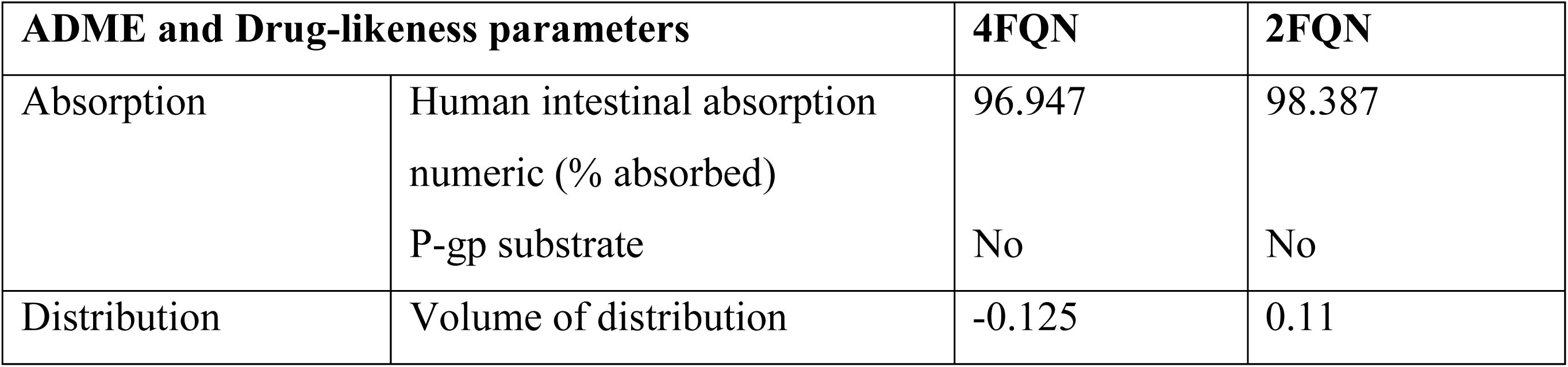
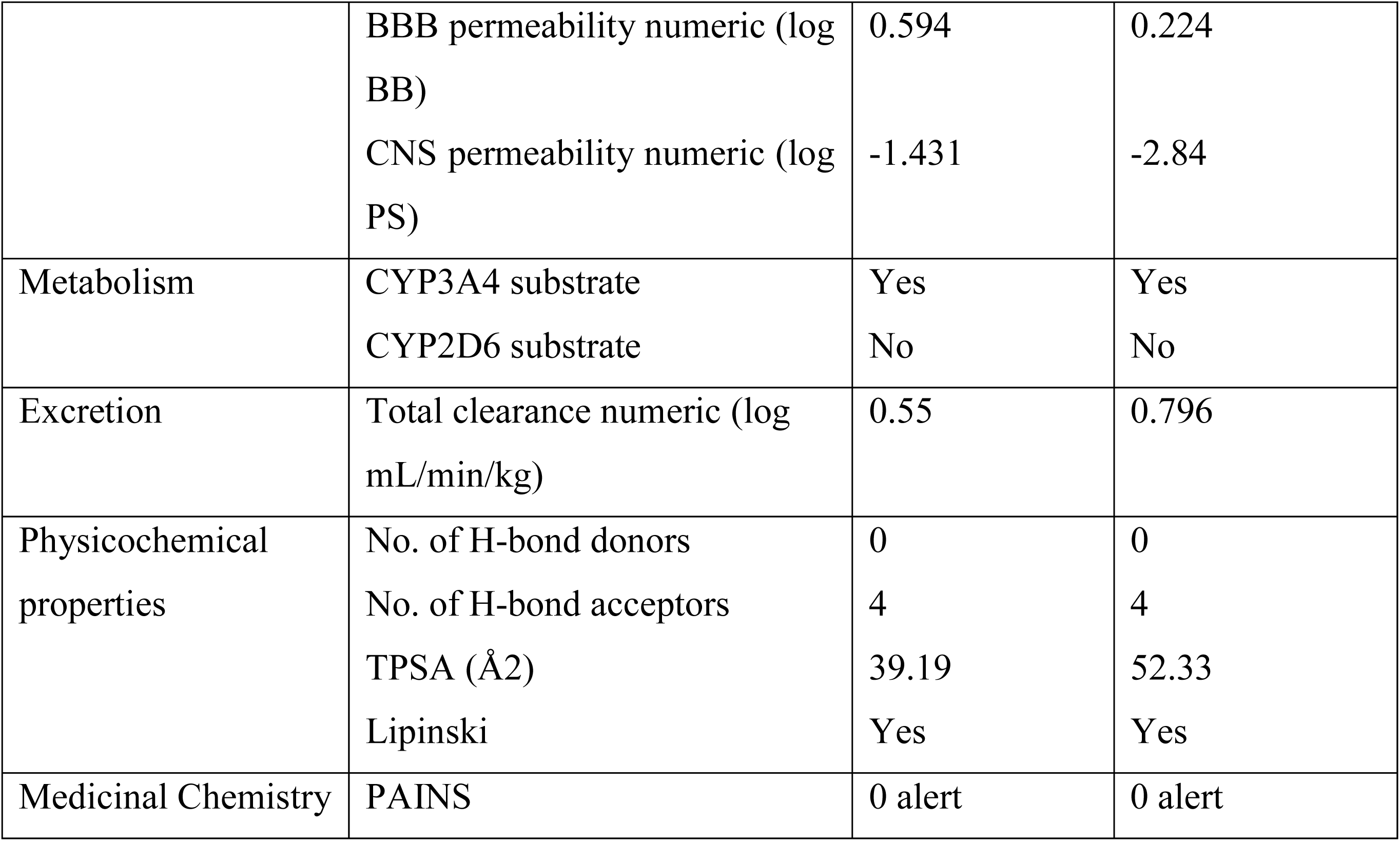
ADME and Drug-likeness properties of compounds 4FQN and 2FQN.

### Molecular Dynamics (MD) simulation analysis

To explore the conformational stability and dynamic behaviour of the protein-ligand complexes, 100 ns molecular dynamics (MD) simulations were conducted using the GROMACS simulation package(Saha *et al*, 2025; Kumar *et al*, 2025). Four ligands, CBR-5992 (reference), 64-9D (parent), 4FQN (most active), and 2FQN (second most active) were simulated in complex with NDH-2 (PDB ID: 9MQZ). All systems were prepared with the CHARMM36 force field using CHARMM-GUI, and the best docking poses served as starting structures. The global stability of the protein was assessed by calculating the root mean square deviation (RMSD) of the protein backbone atoms throughout the simulations. As shown in (Figure SI 6), all complexes exhibited an initial equilibration phase during the first ∼10 ns, after which the trajectories stabilised. The protein backbone RMSD for CBR-5992 averaged 0.44 nm (range 0.27-0.65 nm), indicating moderate structural deviations but overall stable behaviour. The parent ligand complex (64-9D) showed slightly reduced RMSD values averaging 0.40 nm (range 0.21-0.57 nm), while the second most active ligand (2FQN) exhibited the lowest backbone RMSD at 0.36 nm (range 0.26-0.50 nm), reflecting particularly tight conformational maintenance. The most active ligand (4FQN) displayed a somewhat higher RMSD average of 0.58 nm (range 0.29-0.80 nm), consistent with modest structural rearrangements but no unfolding. Ligand RMSD analysis (Figure SI 3). was performed to evaluate ligand positional stability within the active site. Ligands 4FQN and 2FQN showed the lowest ligand RMSD averages at 0.13 nm and 0.10 nm, respectively, indicating tight and stable binding. In contrast, the reference (CBR-5992) and parent (64-9D) ligands displayed higher ligand RMSD values of 0.17 nm and 0.20 nm, respectively, with broader fluctuations. These findings suggest that the active compounds maintain more consistent binding poses within the pocket.

Local flexibility of the protein was assessed using per-residue root mean square fluctuation (RMSF) of backbone atoms (Figure SI 6). All four complexes exhibited broadly similar RMSF profiles with average fluctuations ranging from 0.17 to 0.21 nm, indicating that ligand binding does not significantly alter protein flexibility. Notably, 4FQN showed slightly higher average RMSF values, likely due to increased mobility in surface loops, but no signs of destabilisation were observed. Peaks in RMSF corresponded to flexible loop or terminal regions common across all complexes. Ligand atomic fluctuations (Figure SI 3). were also examined through ligand RMSF analysis, revealing that 4FQN has the lowest internal flexibility, consistent with its tight binding profile. The parent ligand 64-9D showed comparatively higher atomic mobility, while CBR-5992 exhibited moderate fluctuations.

The radius of gyration (Rg) analysis (Figure SI6). demonstrated that all four ligand-protein complexes, CBR-5992, 64-9D, 4FQN, and 2FQN, maintained relatively compact and stable conformations throughout the simulation. Rg values started at approximately 2.30 nm and reached up to 2.45 nm, with an average around 2.35 nm. This narrow fluctuation range indicates the absence of significant unfolding or expansion, suggesting that each ligand remained tightly packed within the protein’s binding cavity. The solvent accessible surface area (SASA) profiles (Figure SI 3). Exhibited moderate fluctuations, ranging from about 210 nm² to 245 nm², with an average value close to 230 nm² across the simulation. These consistent values imply that the ligands stayed predominantly buried within the protein’s binding site, maintaining a balanced network of hydrophobic and hydrophilic interactions with surrounding residues.

Hydrogen bonding analysis (Figure SI 6). revealed distinct interaction patterns among the four complexes. The CBR-5992 complex consistently formed two hydrogen bonds, with occasional transitions to three, reflecting a stable yet moderately flexible interaction. The 64-9D complex showed the highest hydrogen bonding propensity, maintaining three hydrogen bonds steadily and frequently increasing to four, five, or even six during the simulation, suggesting particularly strong and persistent interactions. The 4FQN complex maintained a baseline of two hydrogen bonds with intermittent shifts to three, while the 2FQN complex displayed a similar pattern, with two consistent bonds and transient increases to three or four. These results collectively highlight the stable binding nature of all four ligands, with 64-9D demonstrating the most robust hydrogen bonding profile.

### Effect of carbon sources on efficacy of NDH-2 inhibitors

The mycobacteria under diseased conditions restricted to access to certain nutrients and use a limited pool of carbon sources as energy supplies, mainly in the form of lipids, and probably lactate. Further, the inability of mycobacteria to shift to fermentation for energy generation due to absence of fermentative (NADH-dependent) lactate dehydrogenase, makes the oxidative phosphorylation pathway strictly essential for growth(Kalia *et al*, 2019b). Hence, to explore the metabolic flexibility of *Mtb*, we assessed the carbon source-dependent activity of NDH-2 inhibitors using growth inhibition assay. It was observed that replacement of glycolytic carbon sources with fatty acids led to a marked increase in the inhibitory potency of the NDH-2 inhibitors. for more effective in media supplemented with fatty acids as carbon sources. Under fatty acid-rich conditions, the test molecules exhibited enhanced suppression of bacterial respiration, suggesting a metabolic dependency that amplifies their efficacy. This carbon source-dependent shift highlights the influence of substrate utilization on drug sensitivity (Figure SI 7).

### *In-vivo* efficacy of the NDH-2 inhibitors

Female BALB/c mice (The Jackson Laboratory) were administered with cyclophosphamide intraperitoneally at a dose of 150 mg/kg body weight on two occasions, separated by a 3-day interval. After immune suppression with cyclophosphamide, mice were infected via nasal route by administering 20 µl (10^5^ CFU) of bacterial suspension (*Mtb* H37Ra) adjusted to an OD600nm of 0.5 approximately equals to 10^7^ CFU/ml.(Kumari *et al*, 2024; Huyan *et al*, 2011) To evaluate the *in-vivo* anti-tubercular efficacy of 4FQN and 2FQN, Balb/c female mice were administered both low and high doses of each compound orally. Drug administration through oral route was started 5 days postinfection with a frequency of five times a week for four consecutive weeks. The therapeutic impact was assessed through comparative analysis of bacterial load reduction, and histopathological findings. This section presents the dose-dependent outcomes and highlights the differential potency of the two quinoline derivatives.

The decline in bacterial load was evident across both low and high dose groups, indicating dose-dependent efficacy. Both 4FQN and 2FQN demonstrated potent bactericidal activity at high dose ie. 10 mg/kg body weight and reduced the bacterial load by 1.26 log_10_ and 1.78 log_10_ respectively (Figure 6A & B). Further, at high doses of 4FQN and 2FQN (10 mg/kg body weight) the histopathological studies revealed the recovery in animals with reduced granuloma, necrosis and immune cells infiltration when compared with untreated vehicle control and CBR-5992 (10 mg/kg body weight), a known NDH-2 inhibitor. 2FQN exhibited superior *in-vivo* efficacy compared to 4FQN, likely reflecting advantageous compound properties.

**Figure 6:**
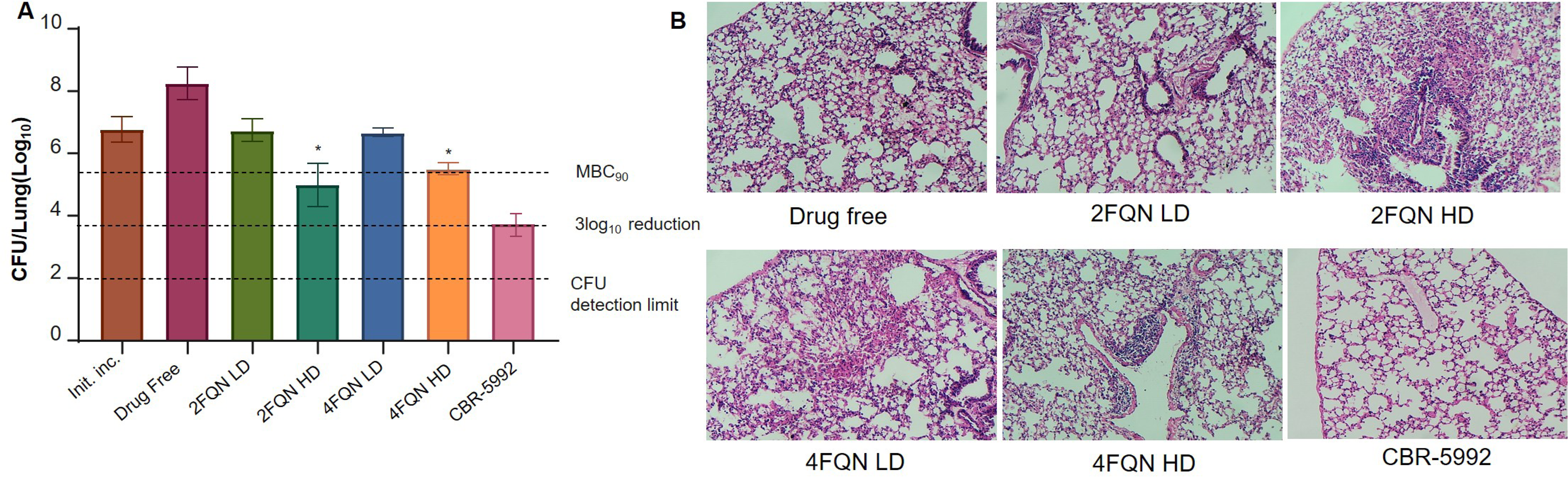
*In-vivo* efficacy of type II NADH inhibitors: BALB/c mice were immunized with cyclophosphamide and aerosol-infected with M. tuberculosis H37Ra. Five days after infection, treatment was started by oral administration of test drugs. Bacterial load (CFU) in lungs of treated animals was assessed after 4 weekk treatment. The efficacy (in form of CFU/lung) of 4FQN-2mg/Kg B.W. (blue bar), 4FQN-10mg/Kg B.W. (Orange bar), 2FQN-2mg/Kg BW (Light green bar), 2FQN-10 mg/Kg BW (Dark green bar) was compared in with initial bacterial load administered (brown bar), drug free control (dark pink bar), and standard NDH-2 inhibitor CBR-5992 (A). Histopathology was performed on all lung samples to determine severity of disease and level of inflammation (B). The data were expressed as mean ± SDs (n=6), with triplicate measurements for each compound concentration. Statistical significance was assessed using one-way ANOVA followed by Dunnett’s test, where \**p* < 0.05, ** *p* < 0.01, and *** *p* < 0.001 compared to the DMSO control.

## Discussion

TB continues to be a serious worldwide health concern, with millions of new cases and deaths each year, highlighting the need for an effective treatment therapy. TB is a major global health concern.

The key objective of this study was to investigate whether inhibition of the NDH-2 enzyme could serve as a viable target for the tuberculosis treatment therapy. Through screening and rational design, we found quinoline-based scaffolds (4FQN and 2FQN) as effective NDH-2 inhibitors that hindered NADH turn over, disrupted ATP homeostasis, and exerted bactericidal action against bith replicating and non-replicating *Mtb*. These molecules have good pharmacokinetic characteristics, long last post antibiotic effect, and higher barrier to resistant development. They also showed specificity for NDH-2 inhibition. Above considering all the results, our results confirm NDH-2 inhibitors as aviable molecular target and highlight the therapeutic benefit of energy metabolism disruption in developing next generation anti-tuberculosis treatment.

The primary screening of a library of 10,000 drug like small molecules was done to identify new scaffolds as probable inhibitors of NDH-2 inhibitors. The shortlisted scaffolds were subjected to molecular modelling using NDH-2 protein structure for protein-ligand interactions and based on binding affinity with the target protein two quinolines based scaffolds were identified as specific inhibitors of NDH-2 i.e. 64-9C and 64-9D. The growth inhibition (MIC) and ATP depletion (ATPIC_50_) values were in correlation and inhibited the mycobacterial growth at very low concentration.

The *in-silico* tools were used to design the structural analogues of 64-9D and total of 10 analogues were synthesized which were evaluated for ATP depletion and growth inhibition using in-vitro assay systems. The optimization of identified hit resulted in enhanced efficacy of the molecules and those which showed better ATP IC_50_ and MIC values were shortlisted for further studies. The *in-silico* binding affinity of all 06 shortlisted molecule was done and the results obtained were compared with the main scaffold and a known NDH-2 inhibitors. The key interactions were observed within the enzyme’s active site, suggesting favorable ligand accommodation. These findings supported their potential as promising NDH-2 inhibitors-with further validation of molecular target using molecular and biochemical assay systems.(Saha *et al*, 2024c)

The importance of NADH in bioenergetics of *Mtb* is very well established as it is an essential electron carrier in the ETC helping to keep membranes energized and maintain ATP homeostasis. Hence, a functional NADH is essential for survival of *Mtb* and minimum NADH turnover is needed. Any, decrease in NADH turnover result in a wider effect on *Mtb*’s overall metabolic activities in addition to influencing the NDH-2 function.(Bhat *et al*, 2016; Yano *et al*, 2011) Therefore, the active molecules were tested for intracellular NADH/NAD⁺ ratio using the peredox-mCherry reporter strain and fluoro NAD kit.(Xu *et al*, 2023a) The results depicted that these inhibitors tremendously impaired the bacterium’s energy metabolism in a dose dependent manner due to their effect on NADH turnover that resulted in reduced ATP levels. The effect of test actives was biochemical assays further confirmed a dose-dependent decrease in NADH oxidation. The inhibition of NADH oxidation in presence of test actives supported our hypothesis and validated these molecules as inhibitors of NADH dehydrogenases.(Saha *et al*, 2025)

To further verify the specificity and effectiveness of the discovered inhibitors, inhibition tests employing purified NDH protein were carried out. All tested molecules demonstrated appreciable inhibition of the NDH enzyme, indicating their potential as effective NDH-targeted agents. Among all probable six test active NDH-2 inhibitors, 4FQN and 2FQN exhibited superior inhibitory activity, outperforming others in the series. Both 4FQN and 2FQN inhibit the NDH enzyme via an un-competitive mechanism, similar to the previously reported mode of inhibition observed for CBR-5992.(Murugesan *et al*, 2018) These findings suggested that the molecules may bind to allosteric sites distinct from the NADH binding pocket, resulting in effective enzyme inhibition independent of substrate concentration.

The inhibition of ATP production by 4FQN and 2FQN in inverted membrane vesicles (IMVs) was observed specifically in the presence of NADH, but not with succinate as the electron donor. This selective inhibition suggested that both compounds target NADH dehydrogenase activity, leaving the succinate-driven respiratory pathway unaffected.(Murugesan *et al*, 2018) These findings further confirmed the specificity of 4FQN and 2FQN towards the NDH-dependent electron transport chain.

Additionally, raising one step mutants, identifying single nucleotide polymorphisms (SNPs) and checking for cross resistance was used as a tool to further validate the NDH-2 as a molecular target for the proposed probable inhibitors.(Samoon *et al*, 2024b; Saha *et al*, 2024c) This experimentation helped in generating important information about genetic variations and mutations that impact NDH-2 function and inhibitor interaction. The identification of SNPs within the coding region of *ndh* gene of one step mutants advanced our knowledge of the function of the enzyme and the selectivity of the inhibitors. These results delineated the role of molecules in inhibiting the function of NDH-2.

Heterogeneity in *Mtb*, posing a serious challenge in anti-TB drug discovery hence, we used a wide range of environmental conditions like replicating, non-replicating (hypoxia and nutrient starved) and macrophage invasion for detailed evaluation of killing efficacy of these NDH-2 inhibitors against different strains of mycobacteria.(Kalia *et al*, 2017a, 2023) Interestingly, these inhibitors effectively eradicated mycobacteria under all the test conditions targeting heterogenous population of mycobacteria. Additionally, the test molecules 4FQN and 2FQN also exhibited time and concentration dependent bactericidal activity against replicating *Mtb* mc² 6230. Notably, complete bacterial killing was observed at 4×MIC on day 5 and similar results were obtained at 2×MIC after exposing the bacteria for 10 days. These results indicated significant antimicrobial action with potential for dose optimization. Further, ATP depletion in non-replicating bacteria by both 4FQN and 2FQN highlights their ability to impair oxidative phosphorylation, consistent with NDH inhibition, even under metabolically dormant states.

*Mtb* is a facultative aerobe and any interference in the functioning of ETC may affect the respiration pattern in mycobacteria. Therefore, these molecules were evaluated for their impact on Oxygen consumption of *Mtb* mc² 6230. A methylene blue based qualitative and MitoXpress probe based quantitative assays were designed to monitor the respiration in bacterial cells during treatment with NDH-2 inhibitors.(Shi Lee *et al*) The decolorization of methylene blue indicated a profound disruption of the electron transport chain whereas the oxygen consumption rate (OCR) was calculated using probe lifetime. The oxygen consumption was reduced by >86% respiration when compared with untreated cells of *Mtb* and this further supported our hypothesis of blocking electron transfer across the ETC resulting in reduced respiration rate followed by abrupt ATP depletion.

Post-antibiotic effect (PAE) evaluation against *Mtb* mc² 6230 revealed that the test compounds exerted prolonged growth suppression following drug removal. Notably, 4FQN exhibited a significantly enhanced PAE of 96 hours at 4×MIC, hence outperformed 2FQN. These findings indicated that 4FQN may exert longer-lasting intracellular activity. The mutation prevention concentrations (MPC) were determined and results revealed the distinct potencies between the tested compounds. The 4FQN completely inhibited the *Mtb* mc² 6230 growth at 16×MIC whereas 2FQN did the same at 32×MIC. Hence the mutation frequency results suggested that both compounds possess a relatively high barrier to resistance development, with 4FQN exhibiting a more robust profile in preventing mutant selection. Combination studies of 4FQN and 2FQN with the electron transport chain inhibitor Telacebec and BDQ using checkerboard assays against *Mtb* mc² 6230 revealed an enhanced antimicrobial response. Both molecules demonstrated interaction profiles approaching borderline synergism, showing greater efficacy than simple additivity.

The *in-silico* ADME profiling of 4FQN, 2FQN, and CBR-5992 indicated favorable oral absorption (>30%), no interaction with the P-glycoprotein efflux transporter, Limited BBB penetration and CNS exposure (log BB < 0.3; logP <−2). Metabolic analysis predicted that all derivatives are CYP3A4 substrates, necessitating dose consideration to avoid drug–drug interactions. Notably, they were neither substrates of CYP2D6 nor inhibitors of CYP3A4. No compound showed clearance-related liabilities, with predicted elimination via both hepatic and renal routes. The oral bioavailability radar plots for 4FQN and 2FQN indicated favorable drug-likeness, with physicochemical properties well-aligned with optimal oral pharmacokinetics.

The MD simulation results collectively demonstrate that the NDH-2 protein forms stable complexes with both novel ligands (4FQN and 2FQN) and the reference compounds CBR-5992 and 64-9D over 100 ns. The lower RMSD and RMSF values observed for 2FQN and 4FQN, along with their stable radius of gyration and consistent SASA profiles, indicate that these active ligands maintain stable and relatively rigid interactions within the enzyme’s binding site. Furthermore, hydrogen bonding analysis revealed a robust and persistent network of interactions, particularly for 64-9D, supporting strong binding affinity. These findings provide valuable insights into the conformational stability and dynamic behaviour of the protein-ligand complexes and underscore the potential of these ligands as effective NDH-2 inhibitors. Carbon source dependent inhibition revealed that the efficacy of NDH-2 inhibitors is significantly influenced by the metabolic context of Mtb, with fatty acid-rich conditions amplifying respiratory suppression.(Kalia *et al*, 2019a) This substrate-dependent sensitivity underscores the pathogen’s metabolic adaptability and suggests that targeting respiration under specific nutrient environments may enhance therapeutic outcomes. Such insights are crucial for optimizing drug regimens that exploit metabolic vulnerabilities. The *in-vivo* efficacy of 4FQN and 2FQN in Balb/c mice underscores their dose-dependent bactericidal potential against *Mtb*, comparable to the NDH-2 inhibitor CBR-5992. Their impact on bacterial load and tissue pathology supports further development of these test molecules as promising anti-tuberculosis candidates.

In summary, adding NDH-2 inhibitors to combination treatment plans provides a multifaceted approach in treating tuberculosis. The revolutionary potential of these inhibitors is highlighted by their ability to avoid the development of resistance, additivity with ETC inhibitors, and targeting heterogenous mycobacterial population resulting in total bacterial sterilization. Their ability to work with ETC inhibitors make them promising option for improving TB therapy and opening the door to more thorough, efficient, and effective therapeutic approaches in the treatment against tuberculosis.

## Statistical analysis

The data were evaluated as mean ± SDs (n=3). Statistical analyses were done using GraphPad prism 8. Differences in variables between were assessed through one way ANOVA, followed by multiple comparisons using Dunnett’s test. Additionally, regression analysis was performed in conjunction with correlation assessment. Statistical significance was set at *p* <0.05.

## Conclusion

The identification of quinoline-based NDH-2 inhibitors signify a transformative step in the pursuit of next-generation tuberculosis therapeutics. By selectively inhibiting NDH-2 an essential respiratory enzyme with no host homologue, these molecules offer a highly specific mechanism of action, and contributing to a more targeted treatment approach. Functional studies confirmed that these compounds disrupt crucial metabolic pathways in *Mtb*, as evidenced by significant decreases in NADH turnover, intracellular ATP levels, and OCR, ultimately impairing the bacterium’s energy metabolism and viability.

Among the series, 4FQN and 2FQN consistently outperformed their analogues, emerging as lead candidates due to their superior NDH-2 inhibition, potent bactericidal activity, and broad efficacy across replicating, dormant, and intracellular *Mtb* populations, including equipotent efficacy towards multidrug resistant. These compounds also demonstrated robust post-antibiotic effects, Comprehensive structure-activity relationship (SAR) optimization, supported by *in-silico* docking, resistance harboring mutations in NDH-2, and enzyme inhibition assays, further validated the specificity and potency of these inhibitors. Collectively, these data provide a compelling rationale for advancing 4FQN and 2FQN into preclinical development. in the ongoing fight against tuberculosis, with the potential to address both active and latent forms of the disease and contribute meaningfully to global TB eradication efforts.

Furthermore, when combined with ETC inhibitors such as Q203 and bedaquiline, 4FQN and 2FQN showed borderline synergistic effects, indicating the potential of co-targeting respiratory complexes to enhance treatment efficacy. *In-silico* analysis revealed that 4FQN, 2FQN, and CBR-5992 possess favorable pharmacokinetic and safety profiles, supporting their potential as orally bioavailable agents. Moreover, 2FQN demonstrated superior dose-dependent *in-vivo* bactericidal efficacy and histopathological recovery compared to 4FQN and CBR-5992.

## Materials and Methods

### Biology

*Mtb* H37Ra was obtained from ATCC, USA. *M. bovis* BCG was generously provided by Dr. Sharmistha Banerjee from the University of Hyderabad. The strains *Mtb* mc^2^ 6230, *Mtb* mc^2^ 8243, and *Mtb* mc^2^ 8259 were kindly gifted by Prof. William Jacob Jr. of the Albert Einstein College of Medicine, USA. Bovine serum albumin (BSA) and DMSO were sourced from Sigma-Aldrich (St. Louis, MO, USA). 7H9 Middlebrook broth and 7H10 Middlebrook agar were acquired from BD Difco Laboratories, USA. Isoniazid, rifampicin, and catalase were obtained from Himedia Labs.

### Bacteria growth conditions

The structural analogues were evaluated to determine the MIC against derivative strains of H37Rv, specifically *Mtb* mc^2^ 6230 and *Mtb* mc^2^ 8243 (isoniazid mono-resistant), *Mtb* mc^2^ 8247 (rifampicin mono-resistant), and *Mtb* mc^2^ 8259 (MDR). The bacteria were cultured in 7H9 liquid broth medium, supplemented with 0.2% glycerol, 0.05% Tyloxapol, and 10% Album-Dextrose-Catalase (ADC), with additional supplementation of L-methionine (50 mg/L), L-leucine (50 mg/L), L-arginine (200 mg/L), and D-pantothenic acid (24 mg/L). The cultures were maintained at 37°C in an atmosphere containing 5% CO₂.

### Determination of minimum inhibitory concentration (MIC)

The bacterial inoculum was obtained from the mid-log phase (OD 600nm range of 0.4-0.6) and adjusted to an OD 600nm of 0.005 (1.0×10⁶ CFU/ml) following standard procedures for MIC determination using a U-bottom 96-well plate (Corning). The final inoculum was diluted to an OD 600nm of 0.005 (1.0×10⁶ CFU/ml) and prepared for use. Each well contained 100 µl of media that was serially diluted twice over the test molecules. Subsequently, 100 µl of mycobacterial strains were introduced into each well and incubated for 8-10 days at 37°C. After the incubation period, the plates were visually assessed, and the lowest concentration of the molecule at which no turbidity was observed was identified as the minimum inhibitory concentration (MIC). Isoniazid and rifampicin were used as standard positive controls.(Samoon *et al*, 2024a)

### Intracellular ATP depletion determination against replicating Mycobacteria

The intracellular ATP levels in the bacteria were assessed using the BacTiter-Glo™ Microbial Cell Viability Assay (Promega, USA). Bacterial cultures were harvested from the mid-log phase and adjusted to an OD 600nm of 0.05. The test molecules were added to each well at the appropriate concentration, and 100 µl of bacterial cultures were dispensed into 96-well opaque white flat-bottom microtiter plates (Corning, USA), followed by incubation at 37°C for 24 hours. After incubation, 25 µl of BacTiter-Glo^TM^ reagent was added to each well and further incubated for 12 minutes. Luminescence was measured using the BioTek Citation 5 Multi-Mode Reader, and results were analyzed using GraphPad Prism 8. CBR-5992 and BDQ were used as positive controls in this experiment.(Kalia *et al*, 2017b; Saha *et al*, 2024c)

### Fluorimetric based determination of NADH/NAD^+^ ratio using biosensors

The *Mtb* strain transformed with pMV762-peredox-mCherry was utilized for NADH/NAD⁺ measurement. The *Mtb*-peredox-mCherry strain was cultured in 7H9 medium supplemented with the necessary nutrients. A 200 µl bacterial suspension was dispensed into each well of a 96-well black flat-bottom plate with a clear bottom (Corning, USA), containing varying concentrations of inhibitors. The plates were incubated at 37°C for 24 hours. Spectral measurements were conducted using the BioTek Cytation 5 Multi-Mode Reader, with emissions recorded at 615 nm and 400 nm, and excitation wavelengths set at 400 nm and 587 nm. The NADH/NAD⁺ ratio in bacterial cells was determined by plotting the green/red fluorescence ratio using GraphPad Prism 8. Clofazimine served as a control in this assay. Additionally, the NADH/NAD⁺ ratio was evaluated using the commercially available Fluoro NAD kit (Cell Technology).(Xu *et al*, 2023b)

### *In-silico* approach to find the binding site

Following the synthesis and in vitro evaluation of the quinoline-based ester derivatives, in silico molecular docking was performed to investigate the binding interactions of the two most active compounds (4FQN and 2FQN), parent compound 64-9D and the reference compound CBR-5992 with the target protein. The crystal structure of the *Mycobacterium smegmatis* NDH-2 (PDB ID: 9MQZ) was retrieved from the Protein Data Bank and used as the receptor. All non-essential components, such as co-crystallised ligands and water molecules, were removed prior to docking. Simulations were carried out using AutoDock Vina v1.1.2, with input files comprising the receptor, ligands, and a configuration file specifying the grid centre and dimensions. Default parameters were applied, with exhaustiveness set to 8, the number of binding modes to 10, and an energy range of 3 kcal/mol. Rigid receptor–flexible ligand docking was employed to account for potential conformational changes at the binding interface.

## Determination of half maximal inhibitor concentration (IC_50_)

### Expression and purification of the NDH-2 proteins

To isolate positive clones, the pYUB28b*-Mtb-*NDH-2 construct was initially transformed into *E. coli* Top10 cells and plated on LB-agar solid medium supplemented with 50 μg/ml hygromycin B. Subsequently, the construct was electroporated into the expressing strain *M. smegmatis* mc² 4517 (a generous gift from Professor William R. Jacobs, Albert Einstein College of Medicine, USA) using a Bio-Rad Gene Pulser instrument with the following parameters: R = 1000Ω, V = 2.5 kV, Q = 25 μF. The positive clones were selected by plating on Middlebrook 7H10 agar medium supplemented with 50 μg/ml hygromycin B and kanamycin.(Lu *et al*, 2018; Murugesan *et al*, 2018; Dam *et al*, 2022)

*Mtb*-NDH-2 expression was carried out using an autoinduction medium composed of ZYM-5052, 2 mM MgSO₄, and 0.2× trace elements, supplemented with 0.05% Tween 80 and 50 μg/ml hygromycin B and kanamycin. The culture was incubated for three days at 37°C. Following incubation, cells were subjected to centrifugation at 4000g for 15 minutes, rinsed with 0.1 M potassium phosphate buffer (pH 6.8) containing 1 mM PMSF, and subsequently resuspended in a lysis solution comprising the same buffer with 1 mM PMSF and an EDTA-free complete protease inhibitor cocktail (Roche). The resuspended cells were sonicated on ice and detergent-solubilized at 4°C for two hours using 2% (w/v) Big CHAPs.

After centrifugation of the detergent-solubilized fraction containing His-tagged MtNDH-2 at 20,000g for 30 minutes at 4°C, the sample was incubated with pre-equilibrated TALON resin (Takara) at 4°C for one hour in a binding buffer containing 0.1 M potassium phosphate (pH 6.8), 0.25% Big CHAP, and 10 mM imidazole. Following incubation, *Mtb-*NDH-2 was eluted using a PD-10 column (Cytiva) with an elution buffer comprising 0.1 M potassium phosphate (pH 6.8), 0.25% Big CHAPs, and 150 mM imidazole, after several washing steps with an imidazole gradient. The buffer was subsequently exchanged to 50 mM HEPES (pH 7.0), 300 mM KCl, 20% glycerol, 2 mM MgCl₂, and 0.2% (w/v) Big CHAPS. Protein concentration was determined using the Bradford reagent.

### Activity assay

The specific activity experiment was conducted using a plate format setup. The decrease in NADH concentration was monitored spectrophotometrically at 340 nm, utilizing an NADH extinction coefficient of 6220 cm⁻¹M⁻¹. Absorbance changes were measured using the BioTek Cytation 5 Multi-Mode Reader.(Murugesan *et al*, 2018; Shirude *et al*, 2012)

Activity of purified protein was measured in a coupled assay where NADH is being oxidized and menadione got reduced, as described by Lu et al., 2022.(Lu *et al*, 2022) The standard reaction buffer contained 50 mM HEPES buffer pH 7.5, 0.008% Brij-35 and 25 mM NaCl and. Reaction was monitored at 340 nm by following the depletion of NADH where NADH was used at 200µM concentration. The reaction was initiated by taking purified NDH-2 protein (1µg/ml) and by adding menadione to it. The reaction was monitored spectrophotometrically by following the absorbance changes at 340 nm after 60 min in a Multimode reader. The specific activities were calculated using an extinction coefficient of 6220 M^-1^ cm^-1^ for NADH at 340 nm. In order to calculate Km for the pure enzyme against the two substrates, the assay was performed by varying NADH concentration from 1000µM to 50µM, keeping menadione concentration fixed at 50µM. Similarly, menadione concentration was varied from 200 µM to 3.125 µM, keeping NADH concentration constant at 250 µM.(Shirude *et al*, 2012) Enzyme activities were calculated at each substrate concentration for both NADH and menadione. Finally, steady state kinetic parameters were determined by fitting the data to Michaelis Menten equation using the non-linear function of GraphPad Prism 8.(Shirude *et al*, 2012; Murugesan *et al*, 2018)

### IC_50_ determination

Inhibitor stock solutions were prepared at a concentration of 10 µg/ml in 100% DMSO. Prior to reaction initiation, the compounds were pre-incubated with NDH-2 in the reaction buffer for 10 minutes at room temperature. Menadione was then added to trigger the reaction. A two-fold serial dilution of inhibitors was performed, yielding a range of final concentrations. IC₅₀ values were computed using nonlinear curve fitting via sigmoidal four-parameter dose-response regression in GraphPad Prism 8.(Lu *et al*, 2022; Shirude *et al*, 2012; Dam *et al*, 2022)

### Determination of biochemical competition assays

Competition experiments were performed to determine whether the inhibitors function by competing with menadione or NADH as substrates. The assays were conducted using varying concentrations of menadione (25–200 μM) or NADH (25–200 μM). The inhibition of His-NDH-mediated NADH oxidation was assessed following established protocols, and NADH kinetic oxidation was monitored accordingly. At least three independent competition assays were carried out, and the resulting data were used to generate Lineweaver-Burk plots using GraphPad Prism (version 8).(Murugesan *et al*, 2018; Dam *et al*, 2022)

### Target Confirmation through One-Step Mutant Generation

One-step mutants for the NDH-2 inhibitor CBR-5992 were selected by plating *Mtb* mc² 6230 at a concentration of 10⁹ CFU/mL onto 7H10 agar plates supplemented with 10% OADC and the NDH-2 inhibitor at a concentration equivalent to 8 times the MIC value.(Samoon *et al*, 2024b; Saha *et al*, 2024a) After three weeks of incubation at 37°C, selected colonies were grown to log phase in 7H9 liquid broth supplemented with 10% ADC. A liquid broth microdilution assay was conducted to assess the susceptibility of these mutants to CBR-5992, while cross-resistance was also examined. Analysis of test compounds in relation to SNPs in the NDH-2 expressing gene identified mutants exhibiting resistance to CBR-5992 at levels exceeding eight times the MIC value. (Samoon *et al*, 2024b; Saha *et al*, 2024c)

### Preparation of nutrient starved culture of *Mtb* mc^2^ 6230

Exponentially growing cultures of *Mtb* mc^2^ 6230 were extracted using centrifugation and subsequently washed twice with prewarmed DPBS (Thermo Fisher Scientific) supplemented with Ca²⁺, Mg²⁺, and 0.025% Tween 80. Before conducting drug sensitivity testing, the cell density was adjusted to an OD 600nm of 0.15 and cultured at 37°C for two weeks.(Kalia *et al*, 2017b)

### Determination of minimum bactericidal concentration (MBC) using replicating and nonreplicating Mycobacteria

A 90% reduction in the bacterial population compared to the initial inoculum was defined as MBC₉₀. CLFZ and BDQ were used as positive controls. Bacteria were exposed to varying concentrations of test molecules for 10 days at 37°C. Following incubation, the bacterial cultures were serially diluted tenfold and plated onto 7H10 agar to determine colony-forming units (CFU). The agar plates were then incubated for 3 to 4 weeks. For MBC determination using non-replicating Mycobacteria, the inoculum was incubated for a designated period.(Kalia *et al*, 2017b; Shi Lee *et al*)

### Bactericidal assay under hypoxic conditions

In a T25 flask maintained within an anaerobic jar, mycobacterial inoculums were adjusted to an OD 600nm of 0.01 in 7H9-ADC media without glycerol. Hypoxia was self-generated within three days. Hypoxic mycobacteria were then exposed to test compounds using anaerobic atmosphere bags on 24-well plates. Wearing built-in gloves, 1 ml of hypoxic bacterial culture was added to each well of a 24-well drug-containing plate within the bag. The plates were placed in an anaerobic jar with a fresh anaerobic sachet and incubated at 37°C. After incubation, bacterial cultures were serially diluted and plated onto 7H10-ADC agar, followed by storage at 37°C for 20-25 days to determine colony-forming units (CFU).(Kalia *et al*, 2023)

### *In-vitro* time-kill studies

The bacterial inoculum was obtained from the mid-log phase (OD 600nm range of 0.4-0.6) and adjusted to an OD 600nm of 0.005 for time-kill kinetics assessment of the test molecules. Bacteria were exposed to varying concentrations of compounds for 3, 6, 9, 12, and 15 days at 37°C. Following incubation, cultures were taken from the respective wells, serially diluted tenfold, and plated onto 7H10 agar. The agar plates were then incubated for three to four weeks to determine colony-forming units (CFU). MBC₉₀ was defined as a 90% reduction in the bacterial population compared to the initial inoculum. BDQ and clofazimine were used as positive controls.(Saha *et al*, 2024a, 2024c)

### Determination of intracellular killing efficacy

THP-1 cells were seeded in 24-well plates at a density of 3 × 10⁶ cells per well following treatment with 200 nM phorbol myristate acetate. After a 24-hour differentiation period, the cell monolayers were infected with *M. tuberculosis* H37Ra at a multiplicity of infection of 10 for 60 minutes. Prewarmed full RPMI medium was added, either with or without test medications. BDQ was applied at 0.5 µg/mL, while CBR-5992 was used at 10× MIC. Mycobacterial viability was assessed by colony-forming unit (CFU) analysis on agar plates after five days of infection.(Vilchèze *et al*, 2018; Lamprecht *et al*, 2016)

## Oxygen consumption assay

### Using Methylene blue as an indicator

In the presence of test molecules, 5 ml of mycobacterial culture (OD 600nm:0.3) was transferred into 5.2 ml screw-cap glass vials. After six hours of treatment at 37°C, methylene blue dye was added at a final concentration of 0.001%. The vials were securely sealed and incubated in a hypoxic jar at 37°C. Following 96 hours of incubation, the vials were removed from the incubator, and images were recorded for analysis.(Kalia *et al*, 2017a)

### Using the MitoXpress® probe

The probe was reconstituted in 1 ml of sterile water prior to the experiment. After dilution to an OD 600nm of 0.5, 150 µl of bacterial cultures were aliquoted into each well of a 96-well black-wall, clear-bottom plate for further analysis.(Shi Lee *et al*) Each well received 8 µl of probe following the addition of 1.5 µl of test compounds to the respective wells. After sealing with two drops of high-sensitivity oil, the plate was placed in a Cytation-5 Cell Imaging Multi-Mode Reader (BioTek, USA) and incubated at room temperature for 10 minutes. Fluorescence was monitored for 6 to 8 hours at 37°C, with dual read time-resolved fluorescence (excitation: 380 nm, emission: 650 nm) recorded every 5 minutes using two integration windows: 30-µs delay with 30-µs measurement time, and 30-µs delay with 70-µs measurement time. Fluorescence lifetime was determined following the manufacturer’s instructions, and data analysis was conducted using GraphPad Prism 8.(Lee *et al*, 2019)

### Determination of frequency of resistance

The *Mtb* mc² 6230 culture was harvested during the exponential phase and concentrated via centrifugation to a cell density of approximately 5 × 10⁹ CFU/ml. Drugs were incorporated into 7H10 agar according to the specified testing parameters. A 200 µL aliquot of culture was added to each agar plate, followed by incubation at 37°C for at least 3 to 4 weeks.(Sharma *et al*, 2014)

### Checkerboard assay

The broth chequerboard micro-dilution method, as described by Raghunathan et al.., was utilized to assess whether two molecules could coexist(Kumar *et al*, 2018). The effectiveness of test compounds was evaluated *in-vitro* using the chequerboard approach, in combination with other electron transport chain (ETC) inhibitors, such as BDQ (ATP synthase inhibitor) and Q203 (cytochrome *bc1* complex inhibitor), as well as first-line anti-TB drugs like isoniazid and rifampicin.

Seven concentrations of the test compounds, ranging from 0.03 to 1.0 μg/ml, were examined. The chequerboard assay was based on minimum inhibitory concentration (MIC) values obtained via the broth micro-dilution method. In microcentrifuge tubes, seven two-fold serial dilutions at 4× the required chemical concentration were prepared. A 50 µl aliquot of each concentration was applied vertically, beginning at the eleventh column of row eight and extending to the second row of the plate. To serve as a drug control, 50 µl of plain media was introduced into the first row of the plate. Each well received 100 µl of inoculum at an OD 600nm of 0.005, followed by incubation for ten days at 37°C. The minimum inhibitory concentration (MIC) was visually determined for each compound individually, in combination with another test compound, and vice versa.(Priya *et al*, 2023)

### Determination of post antibiotic effect (PAE)

To determine the post-antibiotic effect (PAE) of *Mtb* mc^2^ 6230, a time-kill assay is employed to assess bacterial regrowth following transient exposure to an antimicrobial agent(Chan *et al*, 2001). Cultures of *Mtb* mc^2^ 6230 are grown to mid-log phase in Middlebrook 7H9 broth, then a final OD 600nm: 0.005 was exposed to the test antibiotic; typically, at 1×MIC, 2×MIC, and 4×MIC for usually 3 hours. After exposure, the drug is removed through serial dilutions (1:1000) with the fresh media only, and the bacteria were resuspended in fresh medium. These treated cultures, along with untreated controls, are incubated further, and aliquots were taken at regular intervals (e.g., 0, 24, 48, and 72 hours) for CFU determination on 7H10 agar plates. The PAE is calculated as the difference in time required for the treated culture versus the control to increase by 1 log₁₀ CFU/ml, providing insight into the duration of suppressed bacterial growth after drug removal.

### Safety assessment using HepG2 cell line

The cytotoxicity of the test molecules was evaluated using the MTT [3-(4,5-dimethylthiazol-2-yl)-2,5-diphenyltetrazolium bromide] assay in the human hepatocellular carcinoma HepG2 cell line (ATCC HB-8065). Cells were cultured in DMEM glucose medium supplemented with 10% fetal bovine serum (FBS), streptomycin (100 µg/mL), penicillin (75 µg/mL), HEPES (10 mM), and L-glutamine (2 mM) at 37°C. At a density of 1×10⁴ cells per well, they were seeded into a 96-well flat-bottom plate (Tarsons) and incubated for 24 hours at 37°C with 5% CO₂. The test molecules were serially diluted tenfold, ranging from 1000 to 0.48 µg/mL. MTT was dissolved in phosphate-buffered saline at a concentration of 2.5 mg/mL and added to each well. Cell viability was estimated by measuring the absorbance of the reduced formazan dissolved in DMSO at 570 nm using the BioTek Cytation 5 Cell Imaging Multi-Mode Reader. Cytotoxicity was expressed as CC₅₀, the concentration causing a 50% reduction in cell viability, and data were analyzed using GraphPad Prism 8,(Munagala *et al*, 2015) tamoxifen was used as a standard.

### Determination of selectivity index (SI)

Selective index of the test molecules was calculated by the ratio of cytotoxicity (CC_50_) to the biological activity (MIC) of a molecule. For a molecule to be in therapeutic window, SI should be >10.(Kumar *et al*, 2018)

### *In-silico* physicochemical properties and ADME profiling

The pharmacokinetic and drug-likeness properties of the selected derivatives, 4FQN and 2FQN, were evaluated to assess their oral absorption potential, distribution characteristics, metabolism, and elimination behaviour. Predictions were performed using the SwissADME and pkCSM web-based platforms. Parameters analysed included human intestinal absorption, P-glycoprotein interaction, blood-brain barrier penetration, CNS permeability, volume of distribution, cytochrome P450 metabolism, and clearance. Additionally, physicochemical descriptors such as hydrogen bond donors/acceptors, topological polar surface area (TPSA), Lipinski’s Rule of Five compliance, and PAINS alerts were assessed to estimate drug-likeness.

### Molecular dynamics (MD) simulation analysis

Molecular dynamics (MD) simulations were conducted following our previously reported protocol, with additional reference to established literature methods. The simulations were performed in GROMACS 2020 to evaluate the dynamic stability and conformational behaviour of the protein-ligand complexes. The crystal structure of Mycobacterium NDH-2 (PDB ID: 9MQZ) was obtained from the Protein Data Bank, and the docked poses of 4FQN, 2FQN, 64_9D and CBR-5992 selected from the best molecular docking results were used as starting structures. Ligand and protein topologies were prepared via CHARMM-GUI using the CHARMM36 all-atom force field. Each complex was placed in a triclinic box, solvated with TIP3P water, and neutralised by adding Na⁺ and Cl⁻ counterions to achieve an ionic strength of 0.15 M, with ion placement determined using the Monte Carlo method. The systems underwent energy minimization using the steepest descent algorithm, followed by equilibration in two consecutive phases: NVT and NPT ensembles, each for 100 ps. Production MD runs of 100 ns were then carried out for all complexes under periodic boundary conditions, maintaining a temperature of 311.15 K and a pressure of 1 bar. Post-simulation analyses including RMSD, RMSF, radius of gyration (Rg), solvent-accessible surface area (SASA), and hydrogen bond profiles—were performed using GROMACS built-in tools, while visualization and graph generation were completed with QtGrace.

### Carbon source dependent inhibition by NDH-2 inhibitors

*Mtb* mc^2^ 6230 was first grown to mid-log phase in 7H9 liquid media supplemented with 10% ADS supplement, 0.2% glycerol and 0.05% of tween 80 (additionally supplemented with amino acids and vitamins as mentioned above), washed twice in 7H9 base medium supplemented with 0.05% tyloxapol, without carbon sources, and re-suspended in the same medium.(Kalia *et al*, 2019a) The cells were inoculated at an initial OD600 of 0.005 in 7H9 liquid broth medium supplemented with 0.1% fatty acid free BSA, 0.8% NaCl, and glucose (0.2%), glycerol (0.2%), pyruvate (20mM), acetate (0.2%), propionate (0.1%) and cholesterol (0.062mM) as sole carbon source. Bacterial growth was monitored at OD600 over time using an Biotek Cytation 5 multimode spectrophotometer.

### *In-vivo* efficacy of NDH-2 inhibitors against mice model of tuberculosis

Mouse studies were performed in accordance with the Institutional Animal Ethics Committee of National Institute of Pharmaceutical Education and Research (NIPER)-Hyderabad, guidelines.(Kalia *et al*, 2017b) The protocols used in this study were approved by the Institutional Animal Ethics Committee of NIPER-Hyderabad (Protocol# NIP/01/2024/PC/596). Female BALB/c mice (The Jackson Laboratory) were administered with cyclophosphamide intraperitoneally at a dose of 150 mg/kg body weight on two occasions, separated by a 3-day interval. After immune suppression with cyclophosphamide, mice were infected via nasal route by administering 20 µl (10^5^ CFU/ml) of bacterial suspension (*Mtb* H37Ra) adjusted to an OD600 of 0.5 approximately equals to 10^7^ CFU/ml.(Kumari *et al*, 2024; Huyan *et al*, 2011) The animals were segregated based on drug doses. In total 6 groups were there in study and each group was having 7 animals. All the test drugs were formulated in 20% D-α-Tocopherol polyethylene glycol 1000 succinate (TPGS). Two test compounds i.e 2FQN and 4FQN were tested at 2mg/kg and 10 mg/kg of body weight, CBR-5992 was used as positive control and vehicle control was administered with 20% of TPGS. Drug administration through oral route was started 5 days postinfection with a frequency of five times a week for four consecutive weeks. Animals were sacrificed, and infection in the lungs was determined by homogenization of infected lungs followed by CFU determination on agar plates. For pathological analysis and histological staining, lung samples were fixed in 10% (vol/vol) neutral formalin, paraffin embedment, and the tissues were sectioned at 5 μm. Sections were either stained with Hematoxylin & Eosin and images has been obtained at 100X.

### Chemistry

All chemicals were purchased from Sigma-Aldrich, TCI Chemicals, SRL Chemicals, and Avra and used as received. Molychem silica gel (60-120 and 100-200 mesh) was used for column chromatography, and thin-layer chromatography was performed on Merck precoated silica gel 60-F254 plates. All other chemicals and solvents were obtained from commercial sources and purified using standard methods.

### Procedure for the synthesis of compounds (PPQN-2FQN)

An oven-dried round-bottom flask was charged with 8-hydroxyquinoline (1 mmol), the respective aliphatic/aromatic acid (1 mmol), and EDC (1.5 equiv), HOBT (1.5 equiv), TEA (1.5 equiv) and DMAP (0.1 eq) in ACN. The reaction mixture was stirred at rt. for 2 h in air. After completion of the reaction, the reaction mixture was extracted with ethyl acetate (15 mL × 3). The combined organic layers were washed with brine, dried over Na_2_SO_4_, and concentrated in vacuo. The residue was purified by column chromatography on silica gel to afford the desired products.

### Quinolin-8-yl propionate

Compound **PPQN** was obtained as a pale white liquid with a 23% yield. ^1^H NMR (500 MHz, CDCl_3_) δ 8.81 (dd, *J* = 4.5, 1.5 Hz, 1H), 8.18 (dd, *J* = 8.5, 1.5 Hz, 1H), 7.50-7.44 (m, 2H), 7.36 (dd, *J* = 8.5, 1 Hz, 1H), 7.23-7.22 (m, 1H), 1.29 (s, 4H), 0.96-0.86 (m, 1H). ^13^C NMR (125 MHz, CDCl3) δ 152.3, 147.9, 138.3, 136.1, 128.5, 127.7, 121.8, 117.9, 110.1, 29.7, 14.1. HRMS-ESI (ESI-TOF) m/z: [M + H]^+^ calcd for C_12_H_12_NO_2,_ 202.0868; found, 202.0867.

### Quinolin-8-yl butyrate

Compound **BQN** was obtained as a brown liquid with a 27% yield. ^1^H NMR (500 MHz, CDCl_3_) δ 8.80 (dd, *J* = 4.5, 1.5 Hz, 1H), 8.19 (dd, *J* = 8, 1.5 Hz, 1H), 7.50-7.46 (m, 2H), 7.37-7.36 (m, 1H), 7.22 (dd, *J* = 7.5, 1 Hz, 1H), 2.37 (t, *J* = 7.5 Hz, 2H), 1.74-1.66 (m, 2H), 0.70 (t, *J* = 7.5Hz, 3H). ^13^C NMR (125 MHz, CDCl_3_) δ 179.00, 152.24, 147.82, 138.29, 136.21, 128.58, 127.78, 121.79, 117.89, 110.19, 35.77, 18.18, 13.59. HRMS-ESI (ESI-TOF) m/z: [M + H]^+^ calcd for C_13_H_14_NO_2_, 216.1025; found, 216.1028.

### Quinolin-8-yl pivalate

Compound **PQN** was obtained as yellowish-white crystals with 89% yield. ^1^H NMR (500 MHz, CDCl_3_) δ 8.80(dd, *J* = 4.5, 1 Hz, 1H), 8.16 (dd, *J* = 8.5, 1.5 Hz, 1H), 7.48 (t, *J* = 8 Hz, 1H), 7.43 (q, *J* = 4.5Hz, 1H), 7.35-7.34 (m, 1H), 7.24 (dd, *J* = 7.5, 1 Hz, 1H), 1.28 (s, 9H). ^13^C NMR (125 MHz, CDCl_3_) δ 185.2, 152.3, 147.7, 138.1, 136.5, 128.7, 127.8, 121.6, 118.0, 110.9, 38.5, 27.0. HRMS-ESI (ESI-TOF) m/z: [M + H]^+^ calcd for C_14_H_16_NO_2,_ 230.1181; found, 230.1185.

### Quinolin-8-yl cyclopentanecarboxylate

Compound **CPQN** was obtained as a brown liquid with a 30% yield. ^1^H NMR (500 MHz, CDCl_3_) δ 8.92 (d, *J =* 2.5 Hz, 1H), 8.17 (d, *J* = 8 Hz, 1H), 7.72 (d, *J* = 8 Hz, 1H), 7.54 (t, *J* = 7.5 Hz, 1H), 7.46-7.41 (m, 2 Hz, 1H), 3.33-3.26 (m, 1H) 2.29-2.13 (m, 4H) 1.92-1.82 (m, 2H) 1.76-1.70 (m, 2H). ^13^C NMR (125 MHz, CDCl_3_) δ 183.3, 152.3, 147.7, 138.1, 136.5, 128.7, 127.8, 121.6, 118.05, 111.0, 43.7, 29.9, 25.8. HRMS-ESI (ESI-TOF) m/z: [M + H]^+^ calcd for C_15_H_16_NO_2_, 242.1181; found, 242.1187.

### Quinolin-8-yl cyclohexanecarboxylate

Compound **CCQN** was obtained as brown liquid with 57% yield. ^1^H NMR (500 MHz, CDCl_3_) δ 8.79 (d, *J =* 3 Hz, 1H), 8.19 (dd, *J =* 8.5, 1.5 Hz, 1H), 7.50-7.44 (m, 2H), 7.36 (d, *J* = 8 Hz, 1H), 7.23 (dd, *J* = 8, 1.5 Hz, 1H), 2.83-2.76 (m, 1H), 1.98-1.82 (m, 5H), 1.78-1.5 (m, 5H). ^13^C NMR (125 MHz, CDCl_3_) δ 183.0, 152.2, 147.8, 138.2, 136.4, 128.6, 127.8, 121.7, 118.0, 110.7, 43.7, 43.6, 29.9, 25.8. HRMS-ESI (ESI-TOF) m/z: [M + H]^+^ calcd for C_16_H_18_NO_2_, 256.1333; found, 256.1329.

### Quinolin-8-yl 4-methylbenzoate

Compound **4MQN** was obtained as a white powder with 83% yield. ^1^H NMR (600 MHz, CDCl_3_) δ 8.95-8.94 (m, 1H), 8.24 (d, *J* = 8.4 Hz, 2H), 8.20-8.19 (m, 1H), 7.75 (dd, *J* = 7.8, 2.4Hz 1H), 7.59-7.55 (m, 2H), 7.43 (q, *J* = 4.8Hz, 1H), 7.32 (d, *J* = 8.4 Hz, 2H), 2.45 (s, 3H). ^13^C NMR (150 MHz, CDCl_3_) δ 165.6, 150.6, 147.6, 144.3, 141.3, 136.2, 130.6, 129.6, 129.3, 126.8, 126.3, 125.9, 121.9, 121.7, 21.8. HRMS-ESI (ESI-TOF) m/z: [M + H]^+^ calcd for C_17_H_14_NO_2_, 264.1019; found, 264.1017.

### Quinolin-8-yl 4-methoxybenzoate

Compound **MOQN** was obtained as a pale white powder with a 74% yield. ^1^H NMR (500 MHz, CDCl_3_) δ 8.91 (dd, *J* = 9.6, 1.8 Hz, 1H), 8.31-8.29 (m, 2H), 8.20 (dd, *J* = 8.4, 1.8 Hz, 1H), 7.79-7.75 (m, 1H), 7.59 –7.55 (m, 2H), 7.43 (q, J = 4.2 Hz, 1H), 7.02-7.00 (m, 2H), 3.90 (s, 3H). ^13^C NMR (125 MHz, CDCl_3_) δ 165.2, 163.9, 150.6, 147.7, 141.5, 136.1, 132.7, 129.6, 126.3, 125.8, 121.8, 121.8, 121.7, 113.8, 55.5. HRMS-ESI (ESI-TOF) m/z: [M + H]^+^ calcd for C_17_H_14_NO_3_, 280.0974; found, 280.0979.

### Quinolin-8-yl 4-fluorobenzoate

Compound **4FQN** was obtained as pale white crystals with a 70% yield. ^1^H NMR (500 MHz, CDCl_3_) δ 8.92 (dd, *J* = 4.5, 2.0 Hz, 1H), 8.42-8.38 (m, 2H), 8.21 (dd, *J* = 8.5, 2.0 Hz, 1H), 7.80 –7.77 (m, 1H), 7.63-7.58 (m, 2H), 7.45 (q, *J* = 4.5Hz, 1H), 7.25-7.22 (m, 2H). ^13^C NMR (125 MHz, CDCl_3_) δ 166.2 (d, *J* = 253.1 Hz), 164.5, 150.6, 147.9, 141.3, 136.0, 133.2 (d, *J* = 9.4 Hz), 132.1, 129.6, 126.2, 125.8 (d, *J* = 2.9 Hz), 121.8, 115.9 (d, *J* = 22.0 Hz),. ^19^F NMR (470 MHz, CDCl_3_) δ 104.47-104.53 (m, 1F). HRMS-ESI (ESI-TOF) m/z: [M + H]^+^ calcd for C_16_H_11_FNO_2_, 268.0768; found, 268.0770.

### Quinolin-8-yl 4-nitrobenzoate

Compound **4NQN** was obtained as a pale white powder with a 59% yield. ^1^H NMR (500 MHz, CDCl_3_) δ 8.91 (dd, *J* = 4, 1.5Hz, 1H), 8.54 (d, *J* = 9Hz, 2H), 8.41 (d, *J* = 8.5Hz, 2H), 8.26 (dd, *J* = 8.5, 1.5Hz, 1H), 7.84 (dd, *J* = 7.5, 2 Hz, 1H), 7.66-7.61 (m, 2H), 7.49 (q, *J* = 4.5Hz, 1H). ^13^C NMR (125 MHz, CDCl_3_) δ 163.6, 150.9, 150.7, 147.2, 140.9, 136.3, 135.0, 131.7, 129.6, 126.5, 126.2, 123.7, 121.9, 121.4. HRMS-ESI (ESI-TOF) m/z: [M + H]^+^ calcd for C_16_H_11_N_2_O_4_, 295.0724; found, 295.0715.

### Quinolin-8-yl furan-2-carboxylate

Compound **2FQN** was obtained as white powder with 68% yield. ^1^H NMR (500 MHz, CDCl_3_) δ 8.94 (dd, *J* = 4, 1.5 Hz, 1H), 8.22 (dd, *J* = 8.5, 1.5 Hz, 1H), 7.81-7.77 (m, 1H), 7.74-7.74 (m, 1H), 7.62-7.58 (m, 2H), 7.55 (dd, *J* = 3.5, 0.5 Hz, 1H), 7.46 (q, *J* = 4.5 Hz, 1H), 6.65 (q, *J* = 2.0 Hz, 1H). ^13^C NMR (125 MHz, CDCl_3_) δ 157.0, 150.7, 147.1, 146.8, 144.0, 141.2, 136.0, 129.6, 126.2, 126.2, 121.8, 121.6, 119.8, 112.2. HRMS-ESI (ESI-TOF) m/z: [M + H]^+^ calcd for C_14_H_10_NO_3_, 240.0664; found, 240.0661.

## AUTHOR CONTRIBUTIONS

S.S. and R.K have contributed equally in this work. N.P.K. and D.K.S designed research; R.K., P.S. synthesized the structural analogues; S.S., A.R., P.K.A. performed the evaluation studies for synthesized molecules using in vitro and in vivo experiments. S.S., R.K, D.K.S., and N.P.K. analyzed data & wrote the paper; all authors contributed in writing the paper.

## ACKNOWLEDGMENTS

We were grateful to the Indian Council of Medical Research (ICMR), New Delhi, Government of India, for supporting this study by awarding research grant (IIRP-2023-5195). We are thankful to Dr. Santosh Kumar Guru for supporting the study with in-vivo facility. The authors also thankful to NIPER Hyderabad, Department of Pharmaceuticals, Ministry of Chemicals and Fertilizer New Delhi, Government of India for providing necessary infrastructure and research fellowship to S.S.

